# Picking winners in cell-cell collisions: wetting, speed, and contact

**DOI:** 10.1101/2022.05.13.491710

**Authors:** Pedrom Zadeh, Brian A. Camley

**Affiliations:** William H. Miller III Department of Physics & Astronomy, Johns Hopkins University, Baltimore, MD; William H. Miller III Department of Physics & Astronomy and Department of Biophysics, Johns Hopkins University, Baltimore, MD

## Abstract

Groups of eukaryotic cells can coordinate their crawling motion to follow cues more effectively, stay together, or invade new areas. This collective cell migration depends on cell-cell interactions, which are often studied by colliding pairs of cells together. Can the outcome of these collisions be predicted? Recent experiments on trains of colliding epithelial cells suggest that cells with a smaller contact angle to the surface or larger speeds are more likely to maintain their direction (“win”) upon collision. When should we expect shape or speed to correlate with the outcome of a collision? We build a model for two-cell collisions within the phase field approach, which treats cells as deformable objects. We can reproduce the observation that cells with high speed and small contact angles are more likely to win with two different assumptions for how cells interact: (1) velocity-aligning, in which we hypothesize that cells sense their own velocity and align to it over a finite timescale, and (2) front-front contact repolarization, where cells polarize away from cell-cell contact, akin to contact inhibition of locomotion. Surprisingly, though we simulate collisions between cells with widely varying properties, in each case, the probability of a cell winning is completely captured by a single summary variable: its relative speed (in the velocity-aligning model) or its relative contact angle (in the contact repolarization model). Both models are currently consistent with reported experimental results, but they can be distinguished by varying cell contact angle and speed through orthogonal perturbations.

## I. INTRODUCTION

Eukaryotic cells do not just live in isolation, but can function in small clumps, sheets, or complex tissues. Understanding the collective cell migration of these groups of cells is essential to the study of embryonic development, wound healing, and cancer metastasis [1–4]. Groups of cells can have different properties than single cells, including the ability to sense shallower chemical or mechanical gradients [5–13], the ability to amplify cues and develop guided migration over long distances reminiscent of the swarming behavior in insects [14, 15], and an increased efficacy of cancer metastasis [16]. These properties arise from intercellular interactions, including in particular the effect of direct cell-cell contact.

A dramatic and well-studied example of an interaction arising from direct cell-cell contact is “contact inhibition of locomotion” (CIL), first observed decades ago by Abercrombie and Heaysman [17] and later observed in neural crest cells [18] and epithelial cells [19, 20]. In CIL, cells that come in contact with one another retract their local protrusions, repolarize, and subsequently migrate away from contact. CIL is regulated by transmembrane proteins such as cadherins and Eph/ephrins, which regulate the Rho GTPases that ultimately mediate the cell’s protrusive activity [18, 21]. CIL and related properties are essential for the collective chemotaxis of neural crest cells in the developing embryo [5], as well as the spreading of Drosophila hemocytes [22]. CIL has also been shown to promote collective migration in epithelial cells in narrow confinement by establishing cell trains [19].

Contact-based interactions can be affected by the geometry of contact between cells – an example of the broader topic of how confinement and geometry can control collective cell migration [23–25]. For instance, interactions may be asymmetric between the head and tail of a polarized migrating cell [20, 26]. In addition, cells crawling on suspended nanometer-scale fibers, which can have very small amounts of cell-cell contact, may crawl past one another instead of exhibiting CIL [27]. In particular, we are motivated by recent work from the Ladoux group [28], in which two trains of Madin-Darby canine kidney (MDCK) cells collided head-on within narrow confinement. Interestingly, they reported that the train that maintained its direction on collision (“won”) had a leading cell with a smaller contact angle with the substrate [28].

How should we interpret the experimental observations of correlations between collision outcome and geometry? Cell geometry is correlated with cell-substrate adhesion and cell speed [29, 30], which both might separately influence collisions. To understand the role of contact geometry in controlling the outcome of cell collisions, we build a 2-dimensional model for cells within the phase field approach. Here, we study head-on collisions between two cells on adhesive substrates, and we characterize the contact geometry by defining the contact angle as the angle formed by the lamellipodium on the substrate. We control the cell’s contact angle by varying interfacial energies and active forces through cell-substrate adhesion, membrane tension, and protrusion strength. Furthermore, we implement two distinct mechanisms for how cell polarity is influenced by the cell-cell interaction: the velocity-aligning [31, 32] and front-front contact repolarization [27, 33] models, both broadly used to describe collective migration in different contexts. [27, 31–36]. Though the models are fundamentally different, both can produce results nominally consistent with experiment, wherein cells with smaller contact angles and faster speeds are more likely to win. Further analysis, however, could distinguish these two approaches, as we find that within the velocity-aligning model, outcomes are best predicted by difference in speeds between cells, while in the contact repolarization model, outcomes depend most strongly on difference in contact angles. Surprisingly, though we are varying three independent parameters over a broad range of different values, we find in both cases that a single relevant summary parameter predicts the cell-cell collisions well.

## II. MODEL

We build a model for cells within the phase field approach [29, 33–35, 37, 38], which can describe cells with an arbitrary deforming shape. Because the experiments of [28] study cells tightly confined on a microstripe, we simplify our model to two dimensions – a “side view” of the cell [29] (Fig. 1). Our model includes cell self-propulsion, adhesion to a substrate, cell tension, and cellcell repulsion and adhesion. Each cell is given a phase field *ϕ*(**r**,*t*), which is zero outside the cell and one inside the cell, implicitly defining the cell boundary as *ϕ* = 1/2. The evolution of the field for cell *i* is governed by energy minimization and advection of the cell boundary [34, 39, 40],

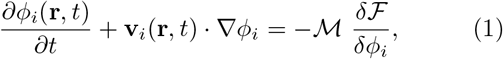

where 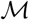 is the transport coefficient and **v**_*i*_(**r**,*t*) is the velocity field of the cell.

**FIG. 1.**
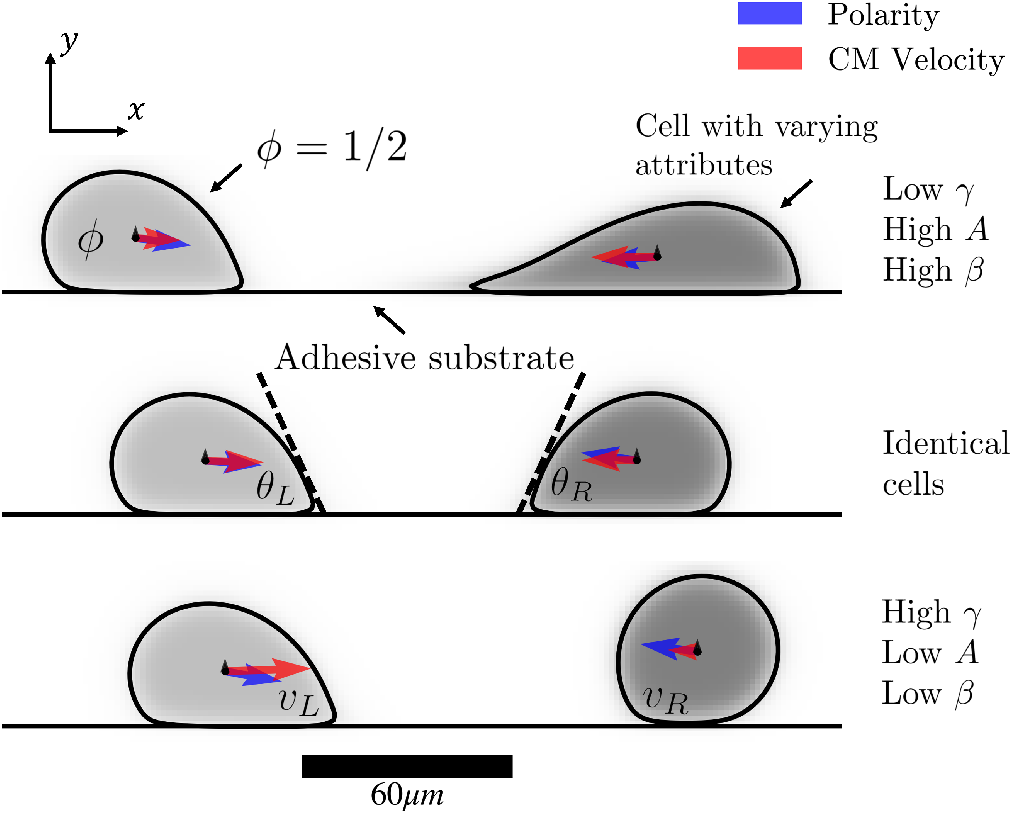
A side-view of two phase field cells wetting on an adhesive substrate and migrating toward each other to undergo a head-on collision. The wide range of accessible contact geometries are shown by varying cell tension *γ*, adhesion *A*, and protrusion strength *β* in the right cell (dark gray): (top) *γ* = 0.9*γ*_0_, *A* = 0.64*γ*_0_, β = 10*γ*_0_, (middle) *γ* = 1.26*γ*_0_, *A* = 0.48*γ*_0_, *β* = 6*γ*_0_, (bottom) *γ* = 1.8*γ*_0_, *A* = 0.32*γ*_0_, *β* = 4*γ*_0_. The left cell (light gray) has constant attributes with the default values listed in Table I. The contact angle of the cell is computed by fitting a line to a few points from the cell’s contour (see Appendix A, Fig. 10b for more details).

The total free energy of the system of *N* cells is 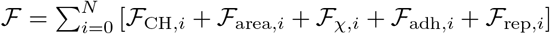. The Cahn-Hilliard energy [34]

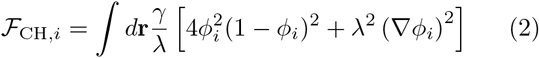

has a double-well potential with minima at *ϕ_i_* = 0, 1 (the outside and inside of the cell) and a gradient term that penalizes interface deformations. *γ* controls the line tension of the cell, and λ has units of length and sets the phase field interface thickness (see Appendix B).

Additionally, we penalize deviations of the cell away from a preferred area 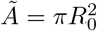 via [34]

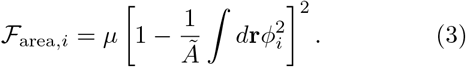

Thus, in the absence of cell-cell interactions, motility forces, and cell-substrate forces, cells relax to circles with radius *R*_0_.

To study lamellipodium contact angles, we must have adhesive substrates onto which cells can wet and extend lamellipodia. We introduce substrates through a static phase field *χ*(*y*) that indicates the substrate, transitioning from zero above the substrate surface to one below the substrate surface (see Appendix A,B). The energy of interaction with the substrate is [29]

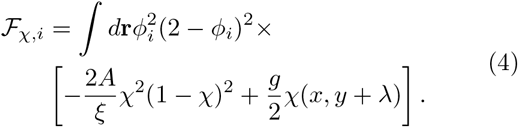

The term proportional to *A* models adhesion. It is nonzero at the boundaries of the cell and the substrate, and it favors increased contact between these two interfaces by reducing the total energy of the cell by an amount proportional to *A*. The term proportional to *g* prevents the cell from penetrating into the substrate. It is nonzero where the boundary of the cell is in contact with the body of the substrate.

Lastly, we account for cell-cell interactions [33, 34, 37]: cells are favored to adhere to each other when their interfaces are in contact,

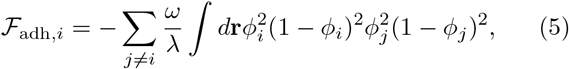

and they are discouraged from overlapping,

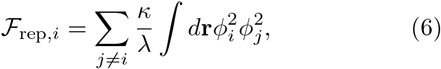

where *ω* and *κ* set the energy scale for each interaction, respectively. We further note that because *ϕ* = 0 is the exterior and *ϕ* = 1 is the interior of the cell, 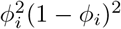 in Eq. 5 is used to indicate the interface of cell *i*.

To complete the phase field description, we obtain the velocity field **v**_*i*_ by noting that cells are overdamped systems. Balancing forces per unit area locally, we write [34]

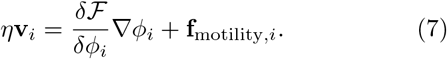

The left hand side represents the friction force per area, while the first term on the right dictates the velocity of the cell boundary driven by energy minimization. The last term, **f**_motility,*i*_, is the motility force per area. The motility force (Section IID) propels the front of the cell forward and depends on cell polarity.

### A. Defining cell polarity

Crawling cells are generally polarized, i.e. they have an asymmetric distribution of proteins and cell shape. For instance, in single cells, Rho GTPase proteins are asymmetric, with Rac1 upregulated in the leading edge of the cell and RhoA active in the cell rear [21]. Rather than explicitly modeling Rho GTPases [33, 35, 41–44], we summarize cell polarity as a single direction *ψ*, with an associated unit vector, 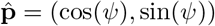.

Choosing how cell polarity reacts to the presence of other cells is an essential stage in modeling collective migration, but there is no single established approach [45]. Here, we implement two separate widely-used alternatives (Fig. 2): a “velocity-aligning” or “self-aligning” polarity, in which cells sense and respond to their own velocity (Section II B), and a “contact repolarization” model, in which cells sense cell-cell contact and repolarize based on that contact (Section II C). We keep the equations of motion between these two approaches as closely analogous as possible.

**FIG. 2.**
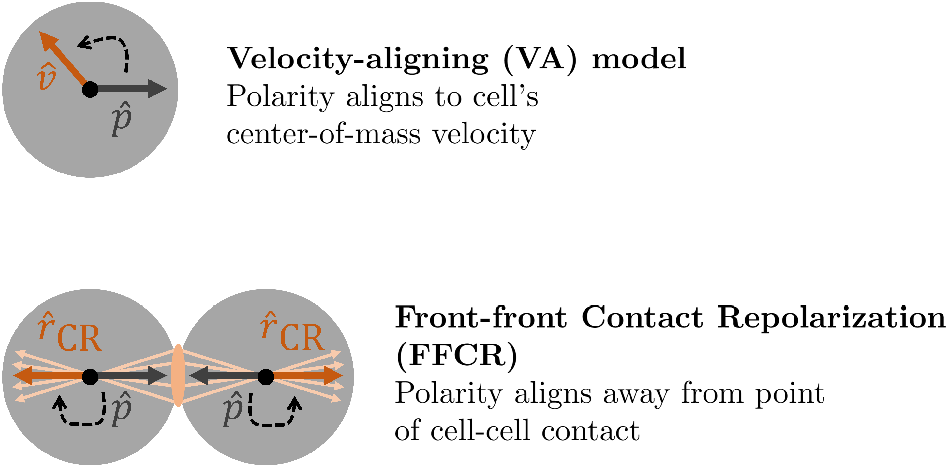
Schematics for velocity-aligning (top) and front-front contact repolarization (bottom) models. In the VA model, cell polarity aligns to the direction of center-of-mass velocity. In the FFCR model, many vectors (light orange) are drawn from the contact region (shaded area) through the center-of-mass of the cell. Cell polarity then aligns to the repolarization vector 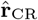, which is calculated as their average (dark orange).

### B. Velocity-aligning cell polarity

The velocity-aligning (VA) model originates from the flocking behavior observed in fish keratocytes [31], and has been extensively used to capture collective migration in different contexts [31–34, 36]. Here, we assume that cells can sense their velocity and repolarize to align to it over a finite timescale *τ*_VA_ (Fig. 2). That is, the evolution of the cell polarity vector is [31, 32]

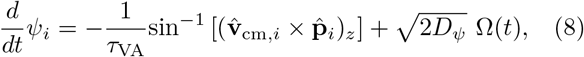

where 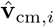 is a unit vector in the direction of cell center-of-mass velocity, *D_ψ_* is the angular diffusion constant, and Ω(*t*) is a Gaussian Langevin noise with 〈Ω(*t*)〉 = 0 and 〈Ω(*t*)Ω(*t*′)〉 = *δ*(**t* – *t*′*). Given this mechanism, the cell polarity integrates information about cell-cell and cell-substrate interactions through the center-of-mass velocity. The inverse sine of the cross product will return the difference between the angle of the center-of-mass velocity and *ψ* when these two angles are close together, but correctly predicts no repolarization when these angles differ by 2*π*. Though each cell only senses its own velocity, because cell-cell collisions lead to correlated velocities, alignment to the cell’s own velocity leads to correlated, coherent migration [31].

### C. Front-front contact repolarization cell polarity

In contact inhibition of locomotion (CIL) [17–20], cells polarize away from cell-cell contact and then migrate away from one another. How do we describe “away from cell-cell contact” in our phase field model? The direction away from a contact point **r**′ is **r**_cm,*i*_ – **r**′, where **r**_cm,*i*_ is the centroid of cell *i*. To define the repolarization direction of cell *i* due to its contact-based interaction with cell *j*, we integrate over all contact points within the contact region *ϕ_i_ϕ_j_* > 0 and constrained to cell *i*. This direction is captured by 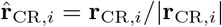 with

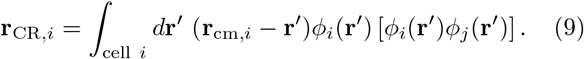

Contact repolarization should only occur when the cells are in contact, which occurs when their phase fields overlap. We thus evaluate Eq. 9 only when max(*ϕ_i_ϕ_j_*) > 0.1 and set **r**_CR,*i*_ = 0 when cell *i* is not in contact with cell *j*. Moreover, we restrict the integral to the region of space where *ϕ_j_* (**r**) > 0.2 to avoid potential minor issues at high wetting when values of *ϕ_j_* outside of the contour *ϕ_j_* = 1/2 may be relevant.

Because we are modeling cells that travel together as a train, cells should no longer repolarize away from contact after one cell has turned around. One possible mechanism is having interactions between cell front and back be different, as supported by [20]. We thus implement a modified version of CIL, front-front contact repolarization (FFCR), in which cells repolarize away from contact only if their fronts are touching [27, 33, 35]. We implement this with a term aligning the polarity of the cell toward the repolarization direction 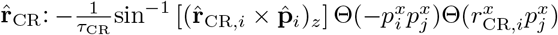, where Θ(*x*) is the Heaviside step function, and 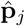 is the polarity of the cell colliding with cell *i*. 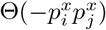 is true when either the fronts of cells *or* the rear of cells are pointing toward each other. To restrict polarity repolarization to only front-front contact, we use 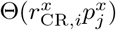, which is true when one cell’s repolarization vector is pointing *away* from the other cell’s front. Together, these two conditions activate contact repolarization only when the fronts of cells are in contact.

With just the contact repolarization term, we would find that the polarity of a single cell would not tend to align along the direction of the substrate, but could point in any direction – this is very different from the VA model. We address this by creating a tendency for the cell to align its polarity along the substrate 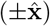 – a “contact guidance” term [21, 27, 46]. In addition, we include an angular noise term Ω(*t*) as in the VA model.

Together, the complete FFCR polarity model is:

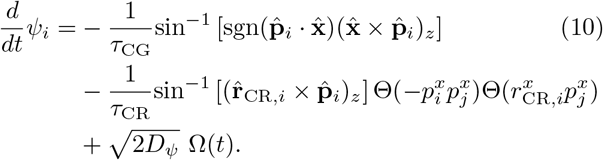

The contact guidance term (the first term) aligns the polarity toward 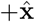 for cells moving to the right and 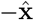 for those moving to the left.

### D. Generating lamellipodium-like protrusions via active forces

We introduce lamellipodia-like protrusions via an active force modeling actin polymerization [21, 47] that is localized to the leading edge of the cell [29, 48]. Mathematically,

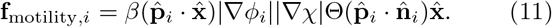

Cells extend lamellipodia on adhesive substrates, and |∇*ϕ_i_*||∇χ| ensures the active force acts only where the cell is in contact with the substrate. To localize protrusions to the leading edge, we compute the normal vector to the boundary, 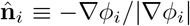, and only allow protrusions when 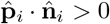. Thus, Eq. 11 yields a motility force that is at the front of the cell, near the substrate, and has a strength proportional to *β* – larger *β* values cause larger protrusions (Fig. 1).

### E. Parameter setting

When possible, the value of a given parameter of the simulation is calibrated to the typical value observed experimentally for trains of MDCK cells confined in narrow channels [28]. Otherwise, values are chosen such that the simulated behavior of cell trains confined in geometries considered in [28] is comparable to what is observed experimentally. The length and time scales of the simulation are chosen such that typical cell sizes and cell speeds are on the order of 40μm and tens of microns per hour, respectively [28].

Line tension *γ* has units of energy/length and sets the energy scale. For living cells, tension consists of mainly two components [49, 50]: (1) membrane tension of the lipid bilayer, and (2) cortical tension from linkage to the actomyosin cortex. We phrase our model so that we do not need to specify *γ*, but only its value relative to some characteristic scale *γ*_0_, which has units of energy/length. All phase field parameters whose units match that of *γ*_0_ are then written as scalar multiples of it: *γ* ~ *γ*_0_, *A* ~ *γ*_0_, *β* ~ *γ*_0_, *ω* ~ *γ*_0_, and *κ* ~ *γ*_0_. The remaining parameters can be written in the following way: 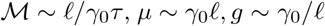, and *η* ~ *γ*_0_*τ*/ℓ, where ℓ and *τ* are our units of length and time (*μ*m and min).

Table I outlines the default values used in the simulation. We set the adhesion strength between cells, *ω*, large enough so that cells with different self-propulsion strengths can still travel cohesively. We set the strength of cell-cell repulsion, *κ*, low enough to ensure cell interfaces can wet against one another to enable intercellular adhesion, yet large enough to prevent cells from overlapping. Strength of repulsion between the cell and the substrate, *g*, is also set following this logic. The constraint penalizing deviations from the preferred area, *μ*, is set relatively strongly to ensure the area varies less than 0.1% from the target area. The alignment time for the VA model is set close to 30 min, the time scale over which the speed of HaCaT cells correlates with their traction forces [51]. In the FFCR model, we choose *τ*_CG_ = *τ*_VA_, so that the single-cell behavior of the two models will be as similar as possible. Lastly, we note that the steady-state target direction in the FFCR model is determined by a competition between the contact guidance and repolarization terms: if contact guidance is too strong (*τ*_CG_ too small), cells will have their polarity strongly constrained to 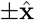 and be unable to repolarize on contact. Solving the deterministic part of Eq. 10, we find that reversals in cell polarity are possible when 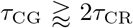 (See Appendix E).

**TABLE I.** Default values of all parameters used in the simulation; any variation from these will be noted. In particular, the values of cell tension *γ*, adhesion to the substrate *A*, and protrusion strength *β* apply to the left cell, whose parameters remain unchanged across all simulations. Numerical integration parameters are given in Appendix D.

| Parameter | Value |
| --- | --- |
| $\gamma$ | $1.26\gamma_0$ |
| $A$ | $0.48\gamma_0$ |
| $\beta$ | $6\gamma_0$ |
| $\omega$ | $30\gamma_0$ |
| $\kappa$ | $1\gamma_0$ |
| $\lambda$ | $4.8 \mu\text{m}$ |
| $R_0$ | $21 \mu\text{m}$ |
| $\xi$ | $3 \mu\text{m}$ |
| $\mu$ | $6000\gamma_0 \mu\text{m}$ |
| $g$ | $8.3\gamma_0/\mu\text{m}$ |
| $\eta$ | $0.67\gamma_0 \text{ min}/\mu\text{m}$ |
| $\mathcal{M}$ | $0.75 \mu\text{m}/(\gamma_0 \text{ min})$ |
| $\tau_{\text{VA}}$ | $24 \text{ min}$ |
| $\tau_{\text{CG}}$ | $24 \text{ min}$ |
| $\tau_{\text{CR}}$ | $12 \text{ min}$ |
| $D_\psi$ | $0.075 \text{ rad}^2/\text{hr}$ |

## III. RESULTS

We simulate collisions between two cells, initialized with a separation distance of 222 *μ*m, migrating toward each other on an adhesive substrate (see numerical methods in Appendix D). In each of these simulations, the left cell has constant parameters (Table I), while the right cell has its properties varied, ranging over (i) tension *γ* ∈ [0.9*γ*_0_, 1.8*γ*_0_], (ii) strength of adhesion to the substrate *A* ∈ [0.32*γ*_0_, 0.64*γ*_0_], and (iii) protrusion strength *β* ∈ [4*γ*_0_, 10*γ*_0_]. Each set contains *n_γ_* = 7, *n_A_* = 11, and *n_β_* = 6 equally spaced points. Together, these form a cubic grid of parameters for the right cell with 462 cell attribute tuples.

We want to see if 1) we reproduce the experimental results showing that winning cells are flatter and faster, and 2) we can understand and predict the outcomes of cell-cell collisions by observing cell properties before collision. We track the contact angle and speed of each cell prior to and during collisions. We also track the “winning probability” *P*_win_, the probability that the cell on the right, whose properties are changing, remains persistent after the collision [28]. To observe this stochastic outcome, we need to run a large number of simulations: we simulate *N* = 96 collisions for each parameter combination (*γ*, *A*, *β*) in the cubic feature-space. We exclude rare simulations where a cell reverses prior to colliding or both cells reverse.

For both models, we found that two characteristics of a collision were crucial: the relative center-of-mass speed *δv* ≡ *v_R_ – v_L_* and the relative contact angle *δθ* ≡ *θ_R_ – θ_L_* (Fig. 1). Here, *v_R,L_* is the speed of the cell on the right (left), averaged over the pre-collision time, and *θ_R,L_* is the contact angle averaged over the pre-collision time. The pre-collision time begins 80 minutes after the start of the simulation, by which point the two cells have equilibrated on the substrate, and ends at the time of collision, where collision is identified with the conditional max(*ϕ_i_ϕ_j_*) > 0.1. Fig. 1 showcases three examples of cell collisions: the top shows a cell pair with *δ_v_* > 0 but *δθ* < 0, the middle shows identical cells, so *δv* = *δθ* = 0, and the bottom has *δθ* > 0 but *δv* < 0.

### A. Relative cell speed controls the winning probability under the VA model

We plot the winning probability as tension *γ*, adhesion *A*, and protrusion strength *β* are varied in the VA model, showing the variation of two parameters at a time, holding the third fixed (Figs. 3A, D, G). Increasing the protrusion strength *β* of a cell increases its probability to win, as does increasing its adhesion to the substrate (Fig. 3A). However, in general, changes in cell tension do not affect the winning probability significantly (Figs. 3D, G).

**FIG. 3.**
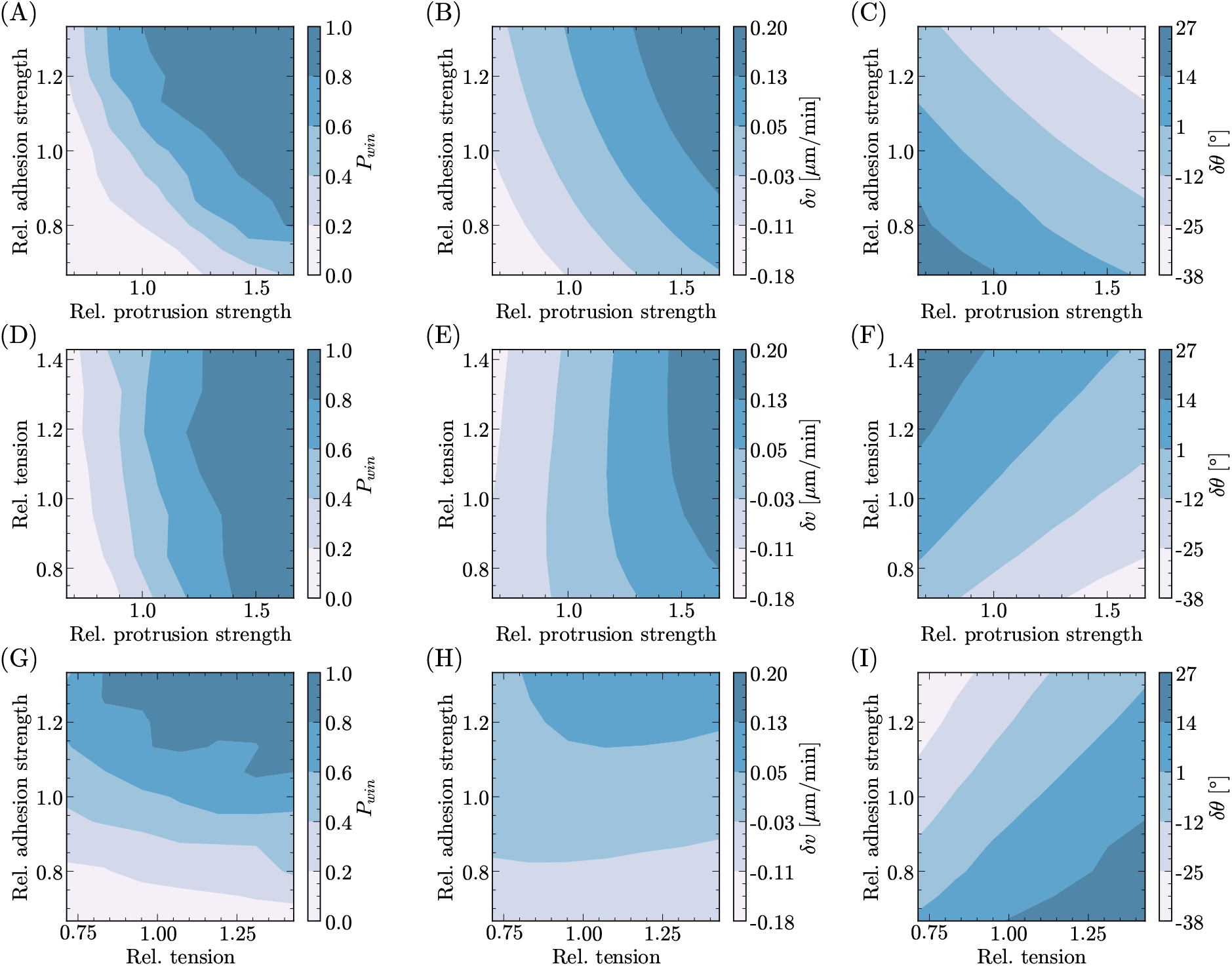
VA model: Winning probability and summary features plotted as a function of two cell attributes with fixed values of (A-C) tension *γ* = 1.2*γ*_0_, (D-F) strength of adhesion to the substrate *A* = 0.48*γ*_0_, and (G-I) strength of protrusion *β* = 6.4*γ*_0_. Contour maps are constructed by running 96 simulations with identical initialization for each parameter set, and the axes are the ratio of the varying cell’s attributes to the left cell’s parameters, which are the default values listed in Table I. Cell polarity is modeled by the velocity-aligning mechanism with *τ*_VA_ = 24 min and *D_ψ_* = 0.075 rad^2^/hr.

The tension *γ*, adhesion *A*, and protrusion strength *β* can also strongly influence the relative speed and contact angle of the two colliding cells. Increasing the right cell’s protrusion strength pushes its front out further, making the cell on the right flatter, leading to negative *δθ*, as does increasing the adhesion to the substrate (Fig. 3C). Meanwhile, increasing the right cell’s tension rounds it up, increasing its contact angle relative to the left cell, making *δθ* positive (Figs. 3F, I). Increasing the protrusion strength increases the speed of the cell, as does an increase in the substrate adhesion (Fig. 3B). The latter effect can be explained by noting that a higher degree of wetting increases the magnitude of |∇_*ϕ*_||∇χ|, and thus results in a higher motility force density. Interestingly, we see that the tension of the cell does not generally affect its speed significantly (Figs. 3E, H).

Are flatter cells or faster cells the winners? When tension is held fixed, the highest winning probabilities correspond to the most negative differences in contact angles (compare top right section of Figs. 3A and C). That is, if we only vary adhesion to the substrate and protrusion strength, we find that flatter cells win more frequently. However, this trend is not true universally: compare *P*_win_ and *δθ* when we vary the cell’s tension (Figs. 3D vs F, 3G vs I). Instead, the pattern that holds true globally is the striking similarity between the distributions of the winning probability and relative cell speed (first and second columns of Fig. 3).

We replot the results of Fig. 3 in Fig. 4, showing how the winning probability depends on *δv* and *δθ* as the parameters *γ*, *A*, and *β* are varied. We find that, as we expected from Fig. 3, winning probability is higher for flatter cells, i.e. *P*_win_ is larger when *δθ* is negative (Fig. 4A). We also see that *P*_win_ correlates well with *δv* (Fig. 4B). Consistent with [28], then, we see that both flatter cells and faster cells are more likely to win. This occurs, however, in part because the speed and contact angle of a cell are highly correlated (Fig. 4C) – cells with larger protrusive strengths in particular are both faster and flatter on average. Comparing Fig. 4A and B, relative speed seems to be a better predictor. Does relative contact angle also predict collision outcomes, or does it simply correlate with relative speed? To address this question, we color the points in Fig. 4 according to which quadrant of the *δv*-*δθ* plot in Fig. 4C they are in. Parameters where *δv* and *δθ* have the same sign are shown in blue and purple, while those where *δv* and *δθ* have different signs are shown in red and green. For the red and green points, the measured contact angle and speed differences predict qualitatively different outcomes. When we look at this subset of parameter values, we no longer observe that flatter cells are more likely to win, but that faster cells are still more likely to win (Figs. 4D, E). In fact, if we perform logistic regressions on *P*_win_(*δv*) and *P*_win_(*δθ*) limited to the subspace where *δv* and *δθ* anticorrelate, we would predict that while faster cells are still more likely to win, cells with smaller contact angles are more likely to *lose*. (We note that here, and in all logistic regressions shown, the regressions are done with scikit-learn’s L2 regularization [52]). This is opposite to what we see in the full parameter set. Based on these results, we expect that – within the VA model of cell polarity – relative cell speed is the essential controlling factor while contact angle merely correlates with speed: when contact angle and speed disagree, we should pay attention to speed. It might seem unavoidable, from our assumptions of the VA model, that cell speed is the controlling variable. However, as we will see in the next section, this is not guaranteed.

**FIG. 4.**
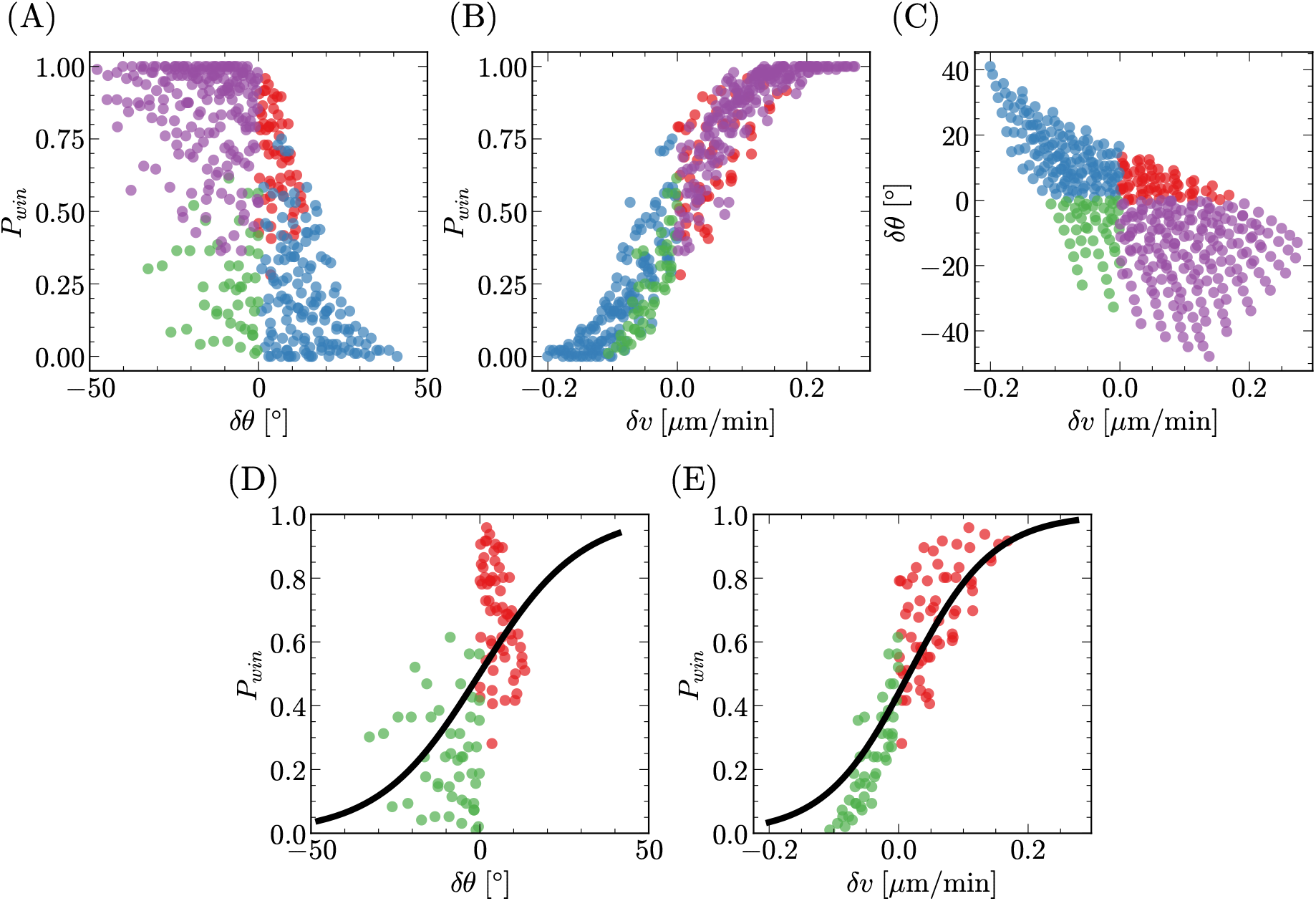
VA model: (A) *P*_win_(*δθ*), (B) *P*_win_(*δv*), and (C) (*δv*,*δθ*) are plotted and colored based on whether *δv* and *δθ* correlate (True: blue, purple; False: green, red). Each scatter point corresponds to different tuples of cell attributes from the cubic feature-space, and it is the average value obtained from 96 simulations. L2-regularized logistic regressions of the form *P*_win_ = [1 + exp(*a*_0_ + *a*_1_*X*)]^−1^ are performed on the subspace in which *δv* and *δθ* do not correlate, where (D) *X* = *δθ* and (E) *X* = *δv*. The training data for these regressions are individual simulations, in which a binary question of *winning* or *losing* is asked.

### B. Relative speed is the sole controlling variable in the VA model only when alignment timescales are long

The alignment timescale *τ*_VA_ plays a large role in controlling collective migration [31, 32, 53], with smaller values of *τ*_VA_ promoting longer, more coherent “trains” of cells in a self-propelled particle model [32]. However, we see a more complex dependence of outcomes on alignment time. In Fig. 5, we repeat the collision simulations for the whole range of parameters *β*, *γ*, and *A* studied in Figs. 3–4, but vary *τ*_VA_ over three values: 80 min, 24 min (our default value), and 4 min.

**FIG. 5.**
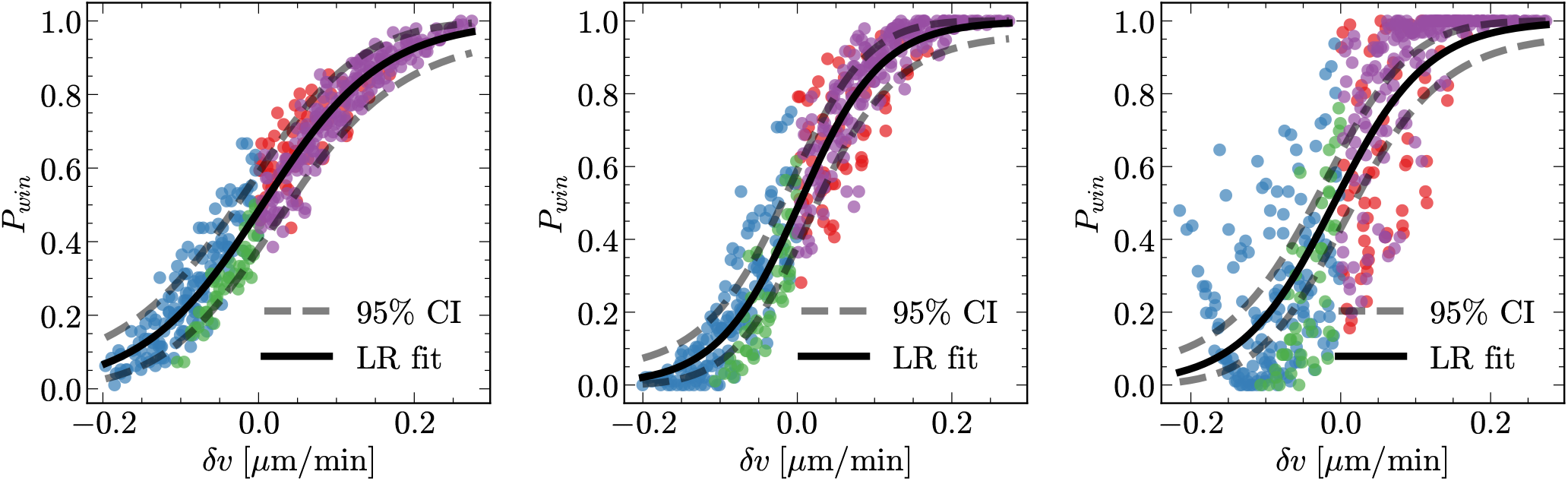
Average winning probability is obtained from 96 simulations with *τ*_VA_ = 80 min (left), 24 min (middle), and 4 min (right). While we observe a sharpening in *P*_win_ as the alignment time decreases, we uncover that at small *τ*_VA_ = 4 min, relative cell speed is no longer predictive of the winning probability, with *P*_win_ scattered far away from the logistic regression fit.

Decreasing *τ*_VA_ suppresses noise in the angle, and makes the cell’s polarity quickly repolarize to its velocity. Our initial expectation about the results of Fig. 5 was that when two cells collided, each cell’s velocity would quickly reach the center-of-mass velocity of the cell pair, which would point in the direction of the faster cell. We would then expect that both cells would tend to polarize in the direction of the faster cell, i.e. the faster cell would win nearly deterministically, predicting that *P*_win_(*δv*) would be essentially a step function. We do see, consistent with this view, that as we decrease *τ*_VA_ from 80 minutes to 24 minutes, the transition becomes sharper and more like a step function (Fig. 5). To some degree this is inevitable – if we took *τ*_VA_ → ∞, each cell polarity’s angle would be an unbiased random walk, and be independent of cell-cell interactions. In this limit, we’d expect *P*_win_ = 1/2 independent of *δv*. However, when *τ*_VA_ = 4 min, instead of a step function in *δv* we see that *P*_win_ no longer collapses neatly as a function of *δv*: there is a huge amount of scatter. This indicates that at small *τ*_VA_, we can no longer reliably summarize all the varying properties of the cells, *A*, *γ*, and *β*, solely by the difference in speeds between the two cells.

The scatter in *P*_win_ is not just due to the finite number of collisions simulated for each point. Suppose *δv* were the sole predictor of the winning probability, such that *P*_win_ = *f*(*δv*), for some function *f*. How large a change in *P*_win_ would we expect to see due to the finite sample size of *n_col_* = 96 collisions? We plot in Fig. 5 the 95% binomial confidence intervals for this binary outcome if *P*_win_(*δv*) were given by the logistic regression fit. (We compute these intervals using the exact Clopper-Pearson method [54].) For the shortest alignment time, *τ*_VA_ = 4 min, the measured *P*_win_ are far more scattered than would be possible if *δv* were a good summary variable. At the longest alignment time we study, *τ*_VA_ = 80 minutes, the quality of the collapse is much stronger, and we are more confident in arguing that *δv* is sufficient to completely predict the outcome of the cell-cell collision.

Why is *δv* no longer a reasonable predictor of the collision outcomes when *τ*_VA_ is small? *δv* is a measure of the difference between cell speeds, averaged over the pre-collision time: it reflects the relative speed in steady state. However, during collision, each cell’s center-of-mass velocity can be altered by local deformations in cell shape. This leads to a transient relative speed during collision, which could be different from *δv* depending on how much each cell deforms. When the alignment time is short, cell polarity integrates velocity information quickly and is affected by instantaneous velocities more strongly – i.e. cell velocity during collision matters more. In this regime, the steady-state relative speed *δv*, which does not properly reflect the transient relative speed present during collision for some simulation parameters, can become a poor predictor of collision outcome. Indeed, when the alignment time is large, cell polarity is driven by cell velocity averaged over that large timescale, and as such, *δv* is able to capture the dynamics of cell collisions more robustly.

Timescales as short as *τ*_VA_ = 4 min are, however, much shorter than current estimates. Recent work reported correlations between the traction forces exerted by Ha-CaT cells and their speed with a time lag of 30 min [51], and interpreted that in terms of a alignment timescale of 30 min.

### C. Relative contact angle controls the winning probability under the FFCR model

We now switch to studying the contact-based FFCR model, and repeat our simulations and analyses from Section IIB using a fundamentally different assumption – that cells repolarize away from front-front contact (Eq. 10).

How do relative cell speed and contact angle predict the winning probability when we vary the parameters *γ*, *β* and *A* in the FFCR model? Given its definition in Eq. 9, the repolarization vector **r**_CR_ is affected by the contact angle of the cell. Cells that extend further on the substrate and form smaller contact angles are also flatter, placing their center-of-mass closer to the substrate and resulting in a repolarization vector closer to the horizontal (Fig. 6). Since contact angle controls the repolarization vector, which is the target direction cell polarity strives to reach, could it also dictate the persistence of the cell and its chances to win?

**FIG. 6.**
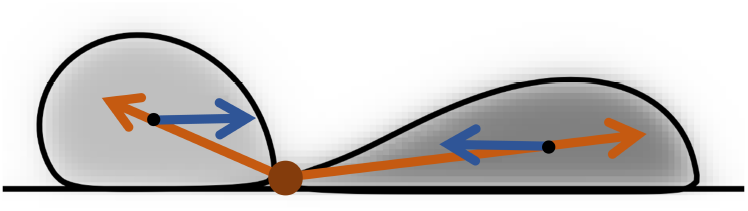
The contact region (dark orange) and contact repolarization vectors 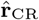 (orange arrows) are drawn for two phase field cells with different shapes. This schematic highlights how the shape of the cell controls the repolarization vector, and how far its polarity (blue arrow) must rotate to reach the desired target direction. While the polarity of flatter cells has a larger angular difference to close, that of rounder cells has a smaller one.

To visualize the outcome of collisions within the cubic feature-space spanned by *γ*, *A*, and *β*, we plot the average winning probability of 96 simulations as a function of two features with the third held constant (Figs. 7A, D, G). Similar to the VA model, we see that increasing the protrusion strength of the cell increases its probability to win, as does increasing its adhesion to the substrate (Fig. 7A). In direct contrast to the VA model, however, we report that increasing the cell’s tension decreases its chances to win (Figs. 7D, G).

**FIG. 7.**
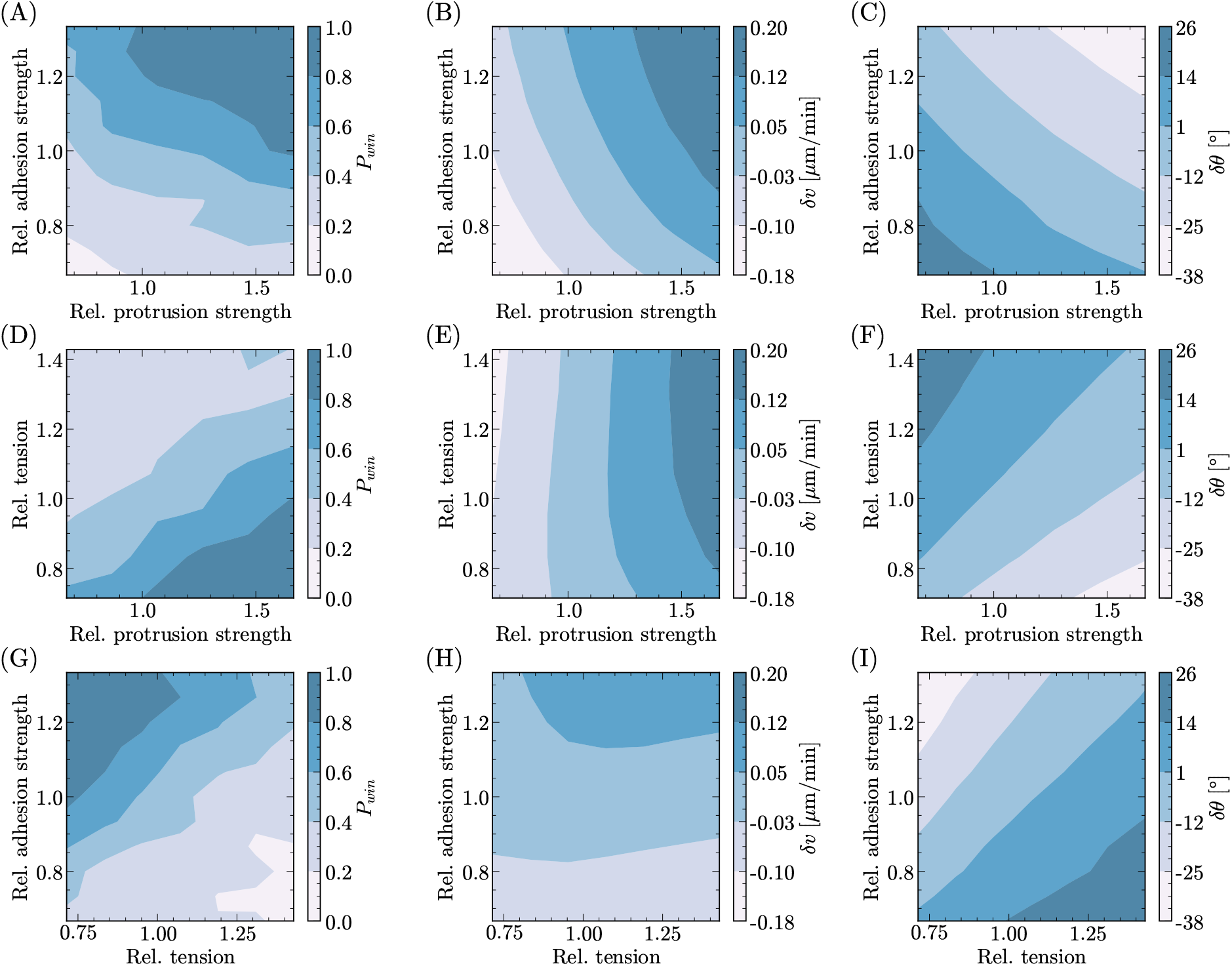
FFCR model: Winning probability and summary features plotted as a function of two cell attributes with fixed values of (A-C) tension *γ* = 1.2*γ*_0_, (D-F) strength of adhesion to the substrate *A* = 0.48*γ*_0_, and (G-I) strength of protrusion *β* = 6.4*γ*_0_. Contour maps are constructed by running 96 simulations with identical initialization for each parameter set, and the axes are the ratio of the varying cell’s attributes to the left cell’s parameters, which are the default values listed in Table I. Cell polarity is modeled by the front-front contact repolarization mechanism with *τ*_CG_ = 24 min, *τ*_CR_ = 12 min, and *D_ψ_* = 0.075 rad^2^/hr.

We also visualize relative speed and contact angle of the two colliding cells as a function of our varied parameters. How the shape and speed of the cell depend on the parameters is highly similar to the results of Fig. 3. Similar to the VA model, we see that increasing the protrusion strength results in flatter and faster cells, as does increasing the adhesion to the substrate (Figs. 7B, C). Moreover, increasing the cell’s tension rounds it up (Figs. 7F, I), while it does not significantly affect its speed (Figs. 7E, H). This correspondence between Fig. 7 and 3 is unsurprising, since *δv* and *δθ* are pre-collision properties. While in principle changing properties of the polarity mechanism can change single-cell properties [32, 33], we chose our two models (Eq. 8 and Eq. 10) to be as close as possible in single cell behavior.

Similar to the VA model, we note that when tension is held fixed, the highest winning probabilities correspond to the most negative differences in contact angles (Figs. 7A and C). In direct contrast to the VA model, however, flatter cells are observed to have higher winning probabilities across the entire feature-space (Figs. 7D and F, G and I). Moreover, the distribution of *P*_win_ is not similar to that of relative speed, but rather to the inverse of relative contact angle (first and third columns of Fig. 7).

We replot the data from Fig. 7 as a function of *δθ* and *δv* in Fig. 8. As in the experiments of [28] and in our earlier VA model, winning probability is generally larger for the faster cell (*δv* > 0, Fig. 8B) as well as for the flatter cell (*δθ* < 0, Fig. 8A). However, in this case it is clear that relative contact angle is the better predictor – the winning probability plotted as a function of *δθ* collapses to a single clear curve, while *P*_win_ only correlates loosely with *δv*.

**FIG. 8.**
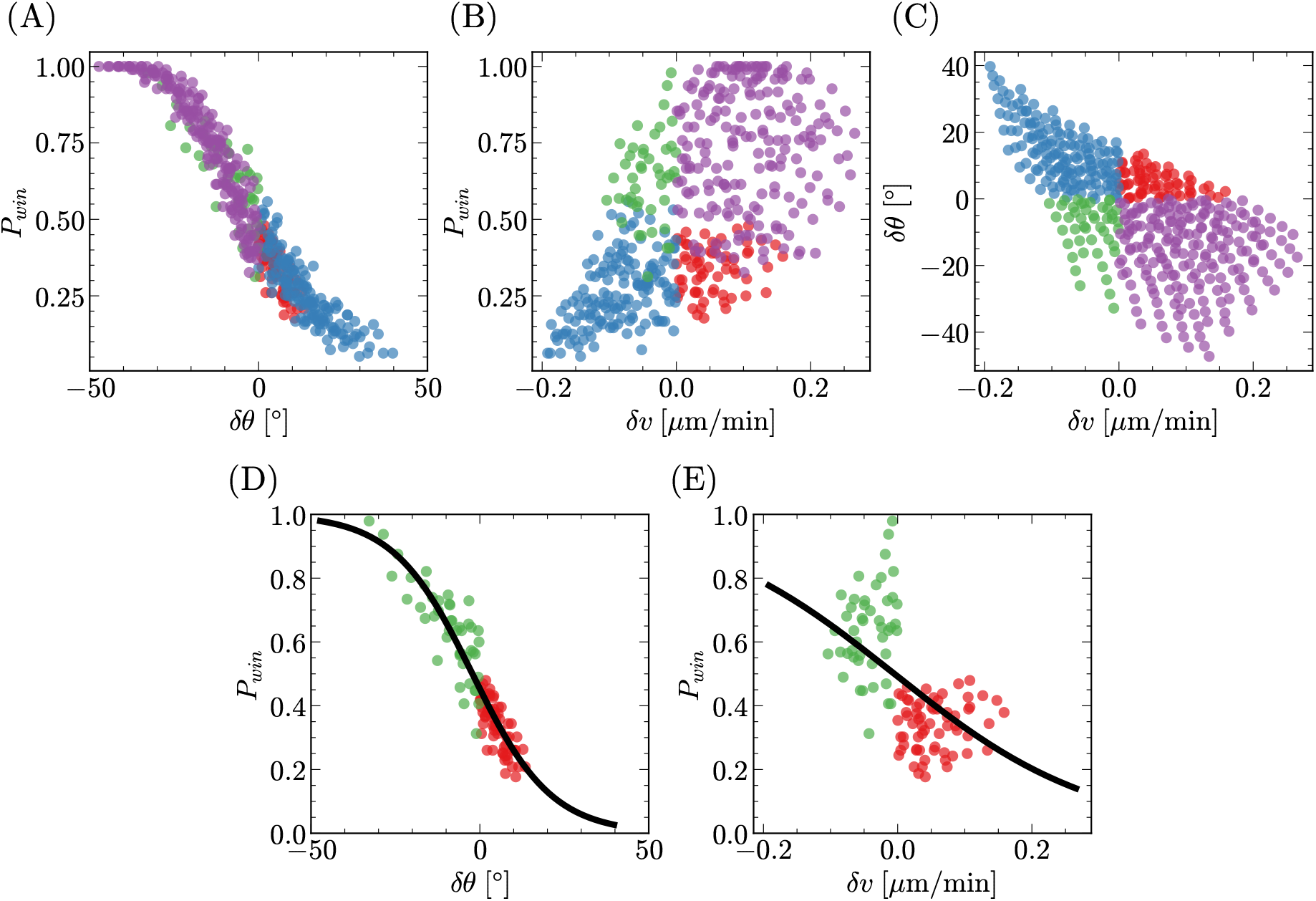
FFCR model: (A) *P*_win_(*δθ*), (B) *P*_win_(*δv*), and (C) (*δv*,*δθ*) are plotted and colored based on whether *δv* and *δθ* correlate (True: blue, purple; False: green, red). Each scatter point corresponds to different tuples of cell attributes from the cubic feature-space, and it is the average value obtained from 96 simulations. L2-regularized logistic regressions of the form *P*_win_ = [1 + exp(*a*_0_ + *a*_1_*X*)]^−1^ are performed on the subspace in which *δv* and *δθ* do not correlate, where (D) *X* = *δθ* and (E) *X* = *δv*. The training data for these regressions are individual simulations, in which a binary question of *winning* or *losing* is asked.

As in Section IIIA above, we explore whether contact angle or speed differences are more relevant in the subspace of parameters where *δθ* and *δv* have opposite signs (red and green points). Within this set of parameters, the winning probability’s dependence on *δv* is opposite to the full space – faster cells are less likely to win when speed and contact angle disagree (Fig. 8E). However, even when speed and contact angle disagree, flatter cells are more likely to win (Fig. 8D). These trends are supported by L2-regularized logistic regressions (black line).

At a surface level, the contact-based FFCR model gives similar predictions to the VA model: faster and flatter cells are more likely to win. However, the underlying driving factor is completely different – cell collision outcomes are completely predicted by the relative contact angle, and the role of speed is only relevant to the extent that it correlates with contact angle.

Why does the FFCR model, Eq. 10, give such a strong dependence on contact angle? At the collision, both cells are being repolarized away from contact – and the cell which turns around first “loses” the collision, as the other cell is no longer in contact with a cell front. We then have to understand why, on average, flatter cells repolarize slower on contact. The FFCR model describes cells repolarizing toward the direction 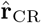, which is strongly influenced by the contact angle of the cells (Fig. 6). Within Eq. 10, the rate of change of cell polarity due to contact repolarization is controlled by a term sin^−1^ 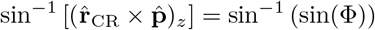, where Φ = *ψ* – *ψ*_CR_ with 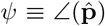 and 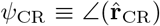 (this term is essentially used to compute the distance between the two angles without giving unphysical results when one angle wraps past 2*π* [31, 32]). sin^−1^ (sin(Φ)) is a sawtooth-shaped graph, which decreases monotonically for Φ ∈ [−*π*, −*π*/2) and increases monotonically for Φ ∈ [−*π*/2, 0]. This means that when the angle from 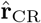 to 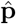 is larger than *π*/2 – i.e. Φ < −*π*/2 – the rate of repolarization decreases as Φ becomes more negative. As an extreme case, if Φ = −*π* and 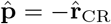, there would be no repolarization – like a pendulum exactly opposed to gravity, this is an unstable equilibrium. For our colliding cells, Φ ∈ (−*π*, −*π*/2). In a typical collision, we see that the flatter cell is repolarized toward a direction 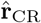 nearly *π* away from its polarity, while the rounder cell has 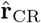 closer to its polarity (Fig. 6). This means that Φ_flatter_ < Φ_rounder_ < −*π*/2. Therefore, the rate of repolarization for flatter cells is smaller and we would expect them to repolarize more slowly and be more likely to win. We can make this argument more explicit by solving a simplified FFCR model (Eq. 10), neglecting stochastic noise and assuming a fixed repolarization direction 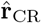. We compute the cell’s repolarization time as a function of the repolarization orientation (see Appendix E). For the most part, this deterministic toy model exhibits a repolarization time that increases with increasing *ψ*_CR_, supporting the notion that flatter cells turn more slowly. However, we do find in some cases, that the deterministic model predicts flatter cells can lose, because of a complicated dependence on the angle *ψ*_CR_ (see Appendix E for a more detailed discussion). We will see in Fig. 9 that, as we make the model more deterministic by decreasing the angular diffusion coefficient *D_ψ_*, contact angle differences do become less predictive of collision outcomes.

**FIG. 9.**
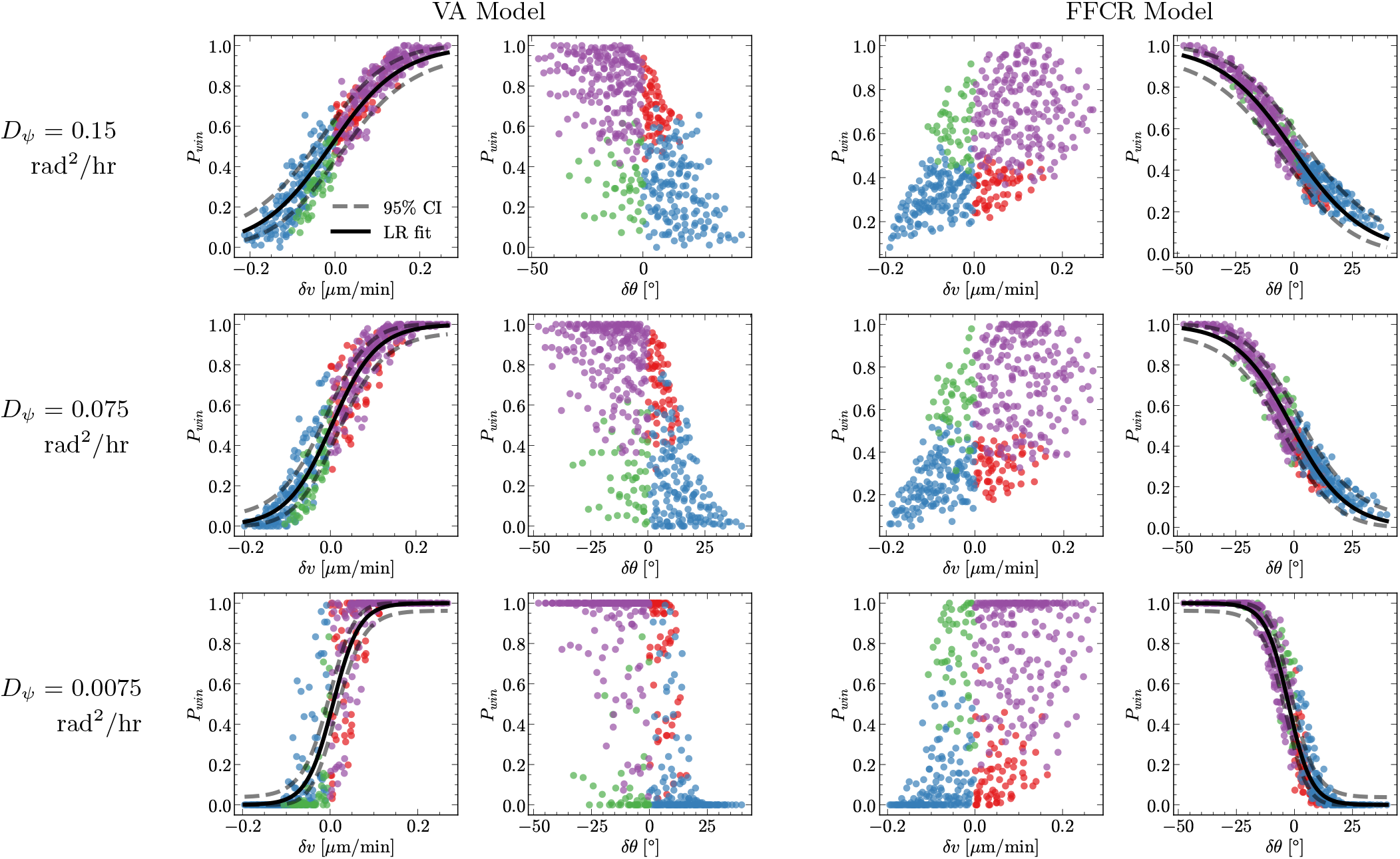
The winning probability plotted against the two summary parameters for different values of *D_ψ_*. L2-regularized logistic regression curves and Binomial confidence intervals are computed for each predictor. We note that lowering the angular diffusion coefficient sharpens the winning probability distribution and increases the scatter of points away from the regression curves.

Our argument above suggests that the outcomes of cellcell collisions can be sensitive not only to cell geometry, but also to the detailed assumptions of the cell polarity model. This makes cell-pair collisions potentially quite a sensitive test of these assumptions.

### D. Angular noise controls how strongly relative speed and contact angle predict *P*_win_

In both the VA and FFCR models, we include an angular Brownian noise: in the absence of polarity mechanisms, the polarity would diffuse with orientational diffusion coefficient *D_ψ_*. These random reorientations model stochasticity arising from finite numbers of molecules in the polarity pathway and from other sources [33, 55–57]. As we have found cell-cell collisions to have stochastic outcomes [19, 28], we would expect this noise, the only random driver in the problem, to play a large role. How does altering *D_ψ_* change outcomes?

We simulated cell collisions at *D_ψ_* = 0.15 rad^2^/hr, 0.075 rad^2^/hr, and 0.0075 rad^2^/hr, with all other parameters kept fixed. Similar to lowering the alignment time, lowering the angular diffusion coefficient makes *P*_win_ a sharper, more step-like function of its relevant predictor, while increasing the scatter of *P*_win_ away from the logistic regression curves (Fig. 9). Similar to our results at small alignment time (Fig. 5), at small *D_ψ_*, these points can be far outside the scatter expected from finite sampling error, as computed by binomial confidence intervals (dashed lines in Fig. 9). As *D_ψ_* decreases – and our model becomes more deterministic – we see that we can no longer summarize the winning probability as a function of a single variable. Again, this indicates that other factors like cell deformability, shape beyond the contact angle, etc., become relevant in this limit.

These results are very similar to those seen in Fig. 5, and likely have a similar origin. We can construct a simple toy model that explains these features qualitatively. A single cell will, because of the angular noise, have a range of possible polarity angles *ψ*. If we linearize the equation of motion of a noncolliding cell’s polarity (Eqs. 8 or 10) around its equilibrium direction (*ψ* = 0 for a cell traveling to the right), we find an Ornstein-Uhlenbeck process, 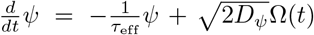, with *τ*_eff_ being *τ*_VA_ or *τ*_CG_ for the VA or FFCR models, respectively. This means that the steady-state probability distribution *P*(*ψ*) for a single cell traveling to the right will be a Gaussian with mean zero and variance proportional to *D_ψ_τ*_eff_ [58]. If this variance is large, then even if the cell on the right has a larger speed than the one on the left on average, at the time of the collision, its speed might be quite different due to the fluctuations in cell polarity. What, then, is the *distribution* of cell velocities entering the collision? Suppose we have two colliding cells with pre-collision average speeds 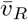 and 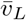. We can model the speeds at collision by adding some fluctuations around the mean: 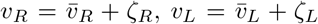. Then, the true difference in speeds at the time of contact will be 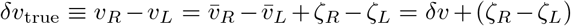, where *δv* is the relative speed averaged over the pre-collision time. Instead of solving the full VA dynamics, we make a simple assumption that the right cell wins when *δv*_true_ > 0 (we can make a similar argument for the FFCR model). With this toy model, the winning probability is then the probability of observing positive true relative speeds: 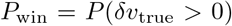. If *ζ_R_* and *ζ_L_* are Gaussian with mean zero and standard deviation *σ*, then 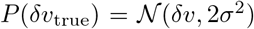. Then, 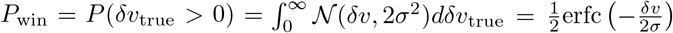, which transitions from 0 to 1 as *δv* moves over a region of order *σ*. If the noise *σ* is small, we would expect even a small change in 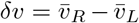 to lead to a large change in *P*_win_. However, if the noise *σ* is large compared to *δv*, then *P*_win_ will not depend much on *δv*. So far, this argument predicts that at low noise levels (small values of *D_ψ_τ*_eff_), we would expect *P*_win_ to become more switch-like and sharper as a function of its relevant variable *δv* or *δθ*. We do see this in Figs. 5 and 9, but we also see that the scatter away from the switch-like curve increases. This can be explained by factors that weaken the correlation between outcome and the predictor – e.g. the deformability at short times or, in case of the FFCR model, the contact angle not correlating with collision outcome in certain cases (see Appendix E). Whatever these factors may be, we can characterize their effect by adding some systematic shift to the relevant variable. For instance, in the VA model, the relevant parameter *δv*_true_ → *δv*_true_ + *ϵ*, and the winning probability becomes 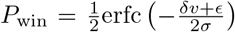. We see that these small, systematic shifts *ϵ* – that differ from parameter set to parameter set – begin to matter more at small values of noise *σ*^2^ ~ *D_ψ_τ*_eff_. Additionally, we would predict that as *D_ψ_τ*_eff_ increases, *P*_win_ becomes more weakly dependent on these shifts. Further increasing *D_ψ_τ*_eff_ to very large levels, we expect *P*_win_ to weakly depend on the predictors and reach a constant value. In this limit, we expect cell polarity to evolve at random and anticipate the outcome of collisions to tend toward 50-50. For instance, the smoothening of the logistic curves we observe in Fig. 5 as the alignment time increases can be explained by the corresponding increase in the variance of *P*(*ψ*).

### E. Predictability of individual cell-cell collisions

Cell-cell collisions are often viewed as entirely stochastic, and models assuming no ability to predict collisions have been successful in understanding some elements of collective migration [19]. However, in our models, we have seen in Figs. 4 and 8 that for a broad range of parameters *β*, *A*, and *γ*, we can reliably predict the probability of outcome by knowing only the relative speed (for the VA model) or only the relative contact angle (for the FFCR model). Suppose we observe a single pair of cells prior to their collision and measure the differences in speeds and contact angles averaged over the pre-collision time, *δv* and *δθ*, respectively. Given this knowledge and our fits above, how reliably can we predict the outcome of this cell-cell collision?

The simplest way to quantify a binary prediction is with a percentage associated with successful labeling of the outcome. We used logistic regression to find a prediction for the winning probability, *P*_win_ = [1 +exp(*a*_0_ + *a*_1_*X*)]^−1^, where *X* = *δv* or *δθ* (see curves in Figs. 4, 8). If, for an individual collision in the VA model, we measure *δv* and predict a win if *P*_win_ > 1/2, what percentage of the time are we correct? We find that (79.5 ± 0.8)% of the time, we can predict the outcome of the VA model with the logistic regression (see Appendix F for details). How does this compare with the simpler – and experimentally accessible – approach of just predicting a win if *δv* > 0? With this criterion, we find that (79.6 ± 0.4)% of the time, we can predict the outcome correctly. Similarly, for the FFCR model, if we use the logistic regression on *δθ*, we can predict the outcome of an individual simulation (72.1 ± 0.7)% of the time, while simply choosing the cell with the smaller contact angle – i.e. *δθ* < 0 – is successful (71.7 ± 0.7)% of the time. We see in both the FFCR and VA models, outcomes are far from a coin flip.

While simply choosing the cell with the faster speed or smaller contact angle to win predicts outcomes well, it doesn’t provide a reliable probability of an outcome, essentially assuming *P*_win_ = Θ(*δv*) for VA and Θ(−*δθ*) for FFCR. A widely used metric for assessing the “goodness of the fit” of a probabilistic binary classifier is the Brier score, the mean squared error between the observed and predicted probabilities, 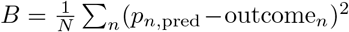, where outcome_*n*_ = 1 for a win and 0 for a loss, and the sum is over all the *N* collisions in the test set (see Appendix F). This penalizes the classifier more if it is overconfident: e.g. for a loss with outcome = 0, *B* is larger when the predicted probability is *p* = 1 than if *p* = 1/2. Lower Brier scores correspond to better classifiers. For the VA model, the logistic regression has a Brier score of 0.140 ± 0.003, while the step-function, assuming *P*_win_ = Θ(*δv*), scores 0.204±0.004. For the FFCR model, the logistic regression has a Brier score of 0.182 ± 0.003, while *P*_win_ = Θ(−*δθ*) scores 0.283 ± 0.007. This signifies that a logistic regression is better at capturing the outcome of simulated collisions with either model than naively choosing the faster or flatter cell.

## IV. DISCUSSION

Cell-cell collisions are broadly used to probe how cells interact and test biochemical regulators of cell interactions [19, 20, 27, 28, 35, 59–62]. To what extent are cell-cell collisions predictable vs purely stochastic? Do cell-cell collisions unavoidably reflect biochemical interactions [18], or do physical properties such as cell shape provide any universal guideline, as is observed in unjamming [63, 64]? How can we use cell-cell collisions to discriminate between competing models of cell interactions? Here, we have focused on answering these questions in a minimal model of cell-pair collisions motivated by experiments of Jain et al. [28], who found that in trains of colliding cells, the winning cell was more likely to have a smaller contact angle and a higher speed. We were able to reproduce these results with a phase field model where the contact angle and cell shape were controlled by tension, adhesion to the substrate, and strength of lamellipodium protrusion. We found initially similar results with two distinct mechanisms for cell polarity – assuming cells align to their own velocity (VA), and assuming cells repolarize away from their front contact region (FFCR). With both assumptions, we were – in certain parameter regimes – able to summarize all the effects of varying adhesion, tension, and protrusion strength in a single variable. This was relative speed *δv* for the VA model and relative contact angle *δθ* for the FFCR model. These predictors were, however, only reliable when collisions were sufficiently stochastic: at small alignment times and small angular diffusion coefficients, cell outcomes reflected changes in parameters outside of this universal response curve.

Our results show that, in order to discriminate between potential polarity mechanisms, it may be essential to study varying multiple cell attributes simultaneously. Our simulations can qualitatively reproduce the results of [28] on cell speed and contact angle with two opposed assumptions, in part because relative cell speed and relative contact angle are correlated in our model. We would be unable to discern whether cell shape is playing an important role in the collision solely based on the result that collision outcome and contact angle are related. To understand this in our modeling, we had to use the multivariable approach of Figs. 4 and 8. Altering parameters like cell-substrate adhesion and protrusion strength simultaneously changed cell speed and contact angle, while altering cell tension allowed us to understand the influence of contact angle independent of speed. Similar approaches could be implemented in cell-cell collision experiments, either via multiple modulations of tension, adhesion, etc., or by exploiting natural cell-to-cell variability. These manipulations might include altering tension via the RhoA-ROCK pathway, adhesion via regulating the concentration of surface proteins or integrin affinity, or lamellipodium protrusion by modulating the activity of membrane-bound Rac [21]. Surface treatment, for instance, has already been seen to regulate contact angle [29, 30]. However, these correlations may be more complex in experimental systems and vary from cell type to cell type. Our minimal model leads to faster cells generally being flatter and more in contact with the substrate; some experiments show shorter cells tending to be faster [65]. Cell-cell collision outcomes may also differ depending on cell type and environment: cell speed and outcome were not noticeably correlated in fibroblast cell-cell collisions on nanofibers [27].

The two polarity mechanisms employed here are relatively simple mathematical caricatures of a complex biological process, but they are commonly employed in modeling collective cell migration [27, 31, 32, 34–36, 66]. Our work shows that these mechanisms can reproduce some essential features of cell-cell collisions. However, in particular for the FFCR model, our results depend not only on the general qualitative structure of the interaction but the specific mathematical details of the mechanism. In particular, we modeled cell polarity as a rigid rotor, such that when given a repolarization cue, it has to traverse a continuum of angles to reach the target direction. The commonly used arcsin approach to handling this sort of cue [31, 32, 67] also controls the outcome, as it ensures stronger responses when the target polarity orientation is closer to the current direction and weaker responses otherwise. These assumptions should ideally be further tested in different contexts. Experimental observations inform us that cell polarity can behave both as an inplane switch and as a rigid rotor, depending on the cell type and context. *Dictyostelium* cells are reported to experience a switch-like, in-plane reorientation of their polarity when the source of chemoattractant cAMP is suddenly moved from facing the front of the cells to the back of the cells [68, 69]. Meanwhile, HL-60 cells and neutrophils have been observed maintaining their polarity and instead executing U-turns to follow the reversal cues [70, 71]. The type of reversal and repolarization can also depend on the amplitude and timescale of the changing signal. The limits of these minimal models can also be better understood by detailed reaction-diffusion modeling of, e.g. Rho GTPases like Rac1 and RhoA on the cell surface [33, 35, 41, 44, 72, 73].

Our result that cell-cell collisions may be predictable, and that varying multiple parameters may be summarized in a single controlling factor, suggests a tantalizing possibility that there may be some degree of universality in cell-cell collision response. To be confident in this idea, experimental tests to see if outcome collisions truly collapse as a function of *δv* or *δθ* are necessary. The key caveat of our results is that – if this universality exists – it is only true to the extent that the underlying repolarization mechanism is conserved.

## ACKNOWLEDGMENTS

We acknowledge support from NIH grant R35GM142847 and NSF grant PHY 1915491. We thank Emiliano Perez Ipiña and Wei Wang for a close reading of the paper and useful comments.

## Appendix A: The phase field model

## Appendix B: The Cahn-Hilliard energy: what do λ and *γ* mean?

The Cahn-Hilliard energy

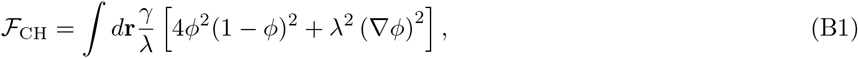

serves to stabilize the outside and inside of the cell and penalizes interface deformations. From dimensional analysis, we conclude that λ has units of length and *γ* has units of energy per length. But what length and line tension are they exactly describing? We briefly derive the energy here; see also [75].

Consider the one dimensional profile of the cell’s phase field taken across a segment perpendicular to its surface (Fig. 10a). Since *ϕ*(*x*) = 0 outside the cell and transitions smoothly to *ϕ*(*x*) = 1 inside the cell, we can postulate the field to be proportional to a sigmoid function around the cell interface: *ϕ*(*x*) = 1/2 + 1/2 tanh((*x* – *x*_0_)/*ξ*), with *ξ* setting the interfacial thickness. To minimize the Cahn-Hilliard energy, this functional form must satisfy the condition 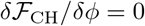. With *f*_CH_ denoting the integrand in Eq. (B1):

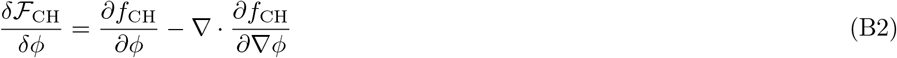

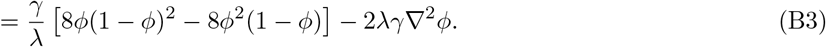

**FIG. 10.**
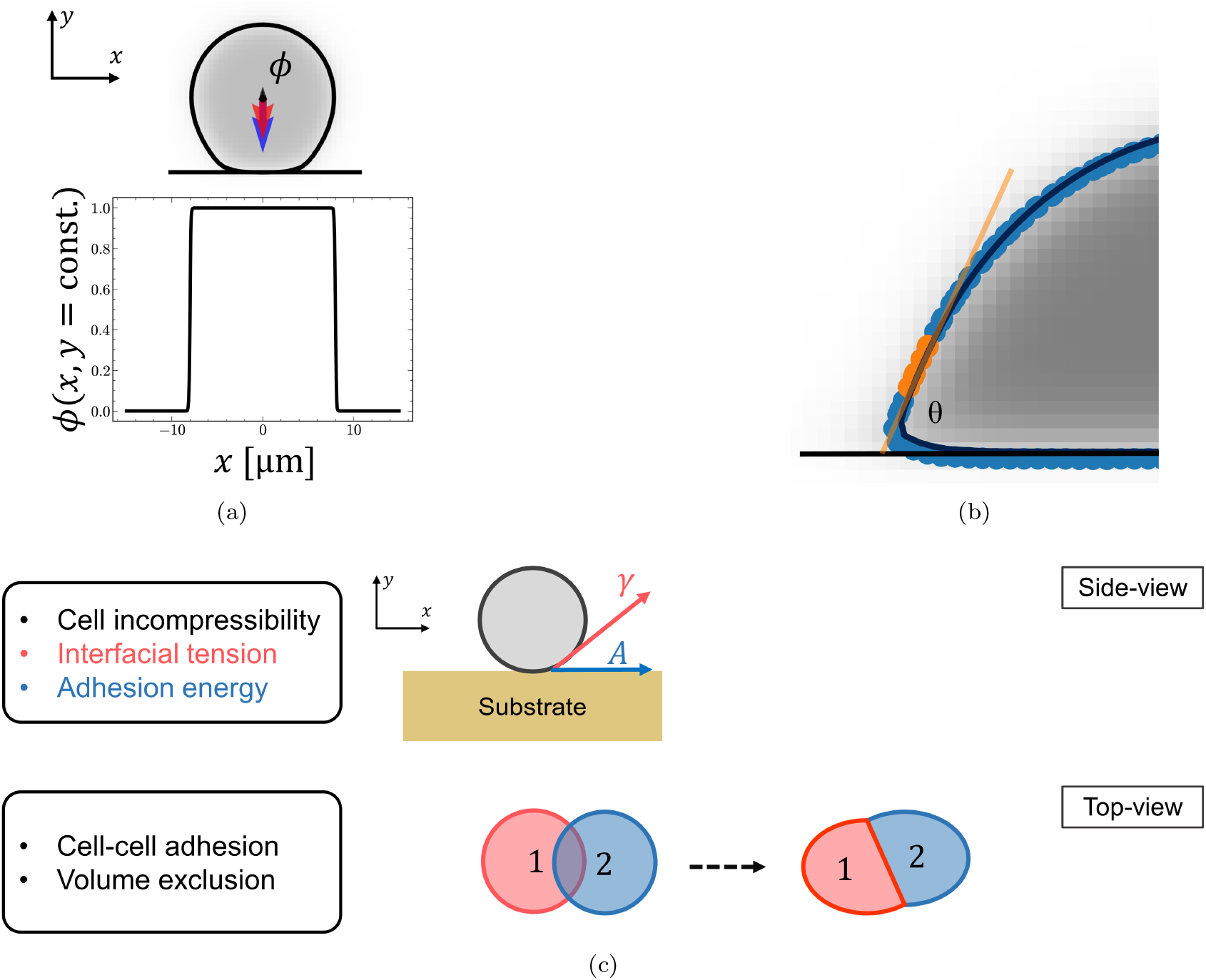
(a) **Elements of the model:** The cell is represented by a smooth, 2-dimensional phase field *ϕ*(**r**,*t*), which is 1 inside the cell and 0 outside the cell. The body of the cell is shown in a grey colormap, and its boundary is drawn in black at *ϕ* = 1/2. The black dot marks the cell’s center-of-mass 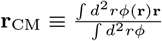, and the arrows depict cell velocity (red) and polarity (blue). The adhesive substrate is defined via the phase field 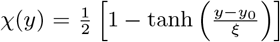, where *y*_0_ defines the location of its interface (*χ*(*y*_0_) = 1/2 drawn in black). Note that 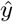 points in the direction above the substrate. The plot below the cell shows the profile of the cell’s field along the *x*-dimension by considering a *y*-slice passing through its center-of-mass. (b) **Contact angle computation:** We compute the contact angle of the cell *θ* formed at the cell-substrate interface by first discretizing the boundary of the cell, located at *ϕ* = 1/2. These contour points (blue circles) are readily obtained from the scikit-image package [74]. A subset of these points, which are appropriately located at the front of the cell (orange circles), are selected and a line is fitted through them. Then, the slope *m* is calculated, and *θ* = arctan *m*. To select the required subset of contour points, we visually inspected the measured contact angle across many simulation parameters and chose the range of points that yielded the most accurate measurements. We then used the same, fixed range across all simulations. (c) **Assumptions of the model:** We model the cell based on the physics of liquid droplets. The cell is incompressible, which in 2D amounts to maintaining a preferred area, and experiences tension across its membrane. It can also wet over an adhesive substrate to lower its energy. Cells interact with each other through cell-cell adhesion, which favors the wetting of their interfaces across one another, and volume exclusion, which penalizes overlap.

Plugging in the sigmoid function, we obtain *ξ* = λ. Thus, we conclude that λ sets the thickness of the cell’s interface.

Moreover, we can compute the total energy of the cell by slicing its boundary into infinitesimally small segments of width *dy*, each of which perpendicular to the cell interface, such that *ϕ*(**r**) = *ϕ*(*x*) for that segment. Then,

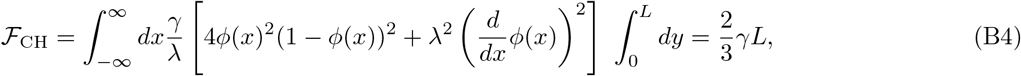

where the integral over *x* computes the energy associated with each segment, and the integral over *y* runs over the total length of the interface *L*. Note that although *ϕ*(*x*) is only valid near the interface of the cell, we can consider the integration bounds {−∞, ∞} for each segment without any issues since the integrand is uniquely zero away from the interface. Notably, we conclude that 2/3*γ* is the line tension of the cell.

## Appendix C: Explicitly writing the phase field equations of motion

The phase field equations of motion for cell *i* are

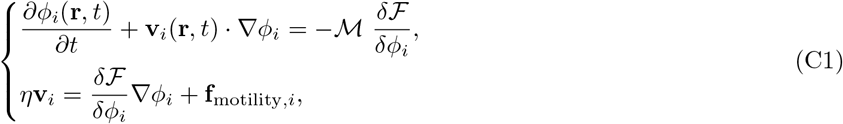

where the total free energy of the system of *N* cells is 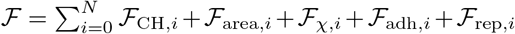. The functional form of each term is given in the main text, and here we will focus on computing 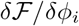 term by term.

1. Focusing on the Cahn-Hilliard energy term:

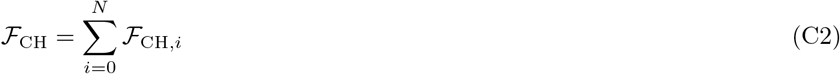

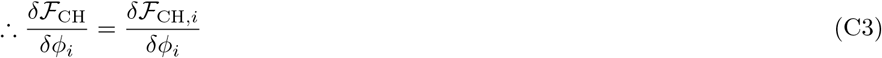

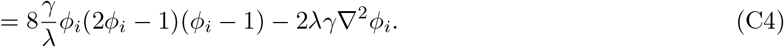
2. Focusing on the area constraint term:

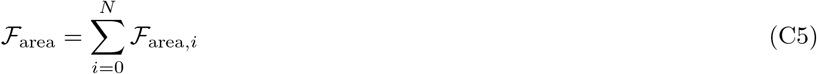

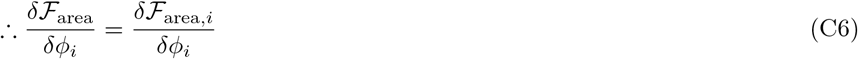

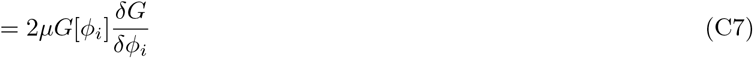

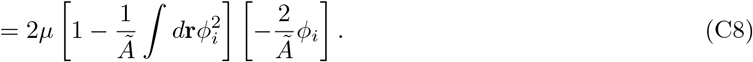 Here, 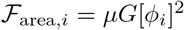, with 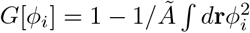.
3. Focusing on the cell-substrate energy term:

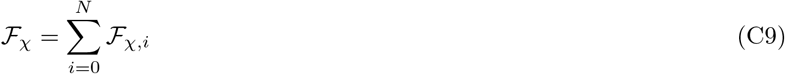

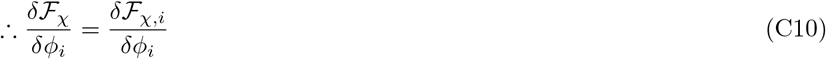

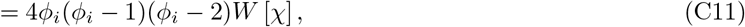

where 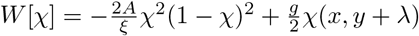.
4. Focusing on the cell-cell adhesion term:

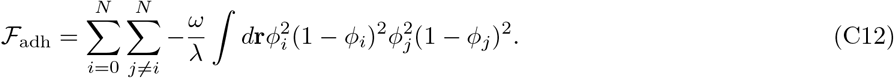 Unlike the previous energy terms, where minimizing the total energy was equivalent to minimizing the energy of the *i*^th^ cell, interaction energies involve cell-pairs. If we expand the terms in Eq. (C12), we would find that the integrand appears *twice*: once when the outer sum is over *i* and once with the outer sum is over *j* ≠ *i* and the inner sum picks *i*. This is due to the integrand being symmetric under *i* → *j*. Thus,

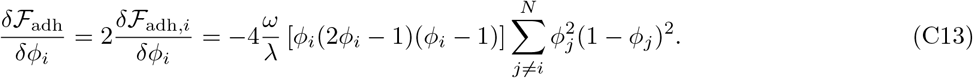
5. Focusing on the cell-cell repulsion term:

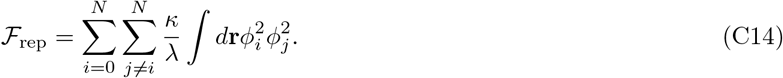 Similar to the cell-cell adhesion energy, the pairwise nature of this term and the symmetry of its integrand under *i* → *j* results in an extra factor of 2. Thus,

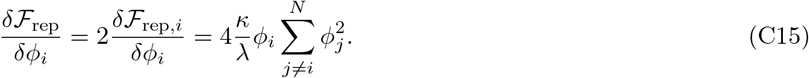

Finally,

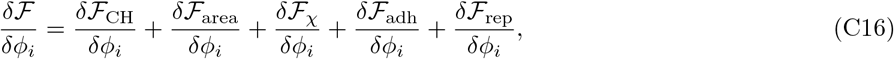

and we have a complete description of the phase field equations of motion (Eq. (C1)) that we can solve numerically.

## Appendix D: Numerical implementation of the model

We numerically solve the phase field equations of motion (Eq. (C1)) within a simulation box of size 300 *μ*m × 300 *μ*m using the explicit finite difference method. With a sufficiently small timestep Δ*t*, the temporal evolution of the phase field is discretized and approximated as

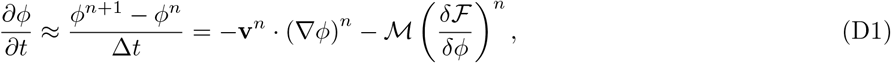

where (·)^*n*^ denotes the value of the quantity at time *t* = *n*Δ*t*. The gradient operator ∇ ≡ (∂_*x*_, ∂*_y_*) is discretized in space at a fixed time as follows, given sufficiently small lattice spacings Δ*x* = Δ*y*:

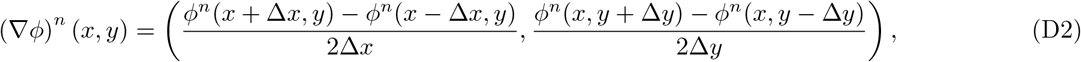

and the Laplacian ∇^2^ ≡ ∂*_xx_* + ∂*_yy_* is discretized using the 4-point nearest neighbor stencil:

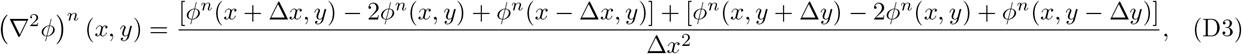

where (∇*ϕ*)^*n*^ (*x, y*) and (∇^2^*ϕ*)*^n^* (*x, y*) are the gradient and Laplacian of the field at time *t* = *n*Δ*t* evaluated at the coordinate (*x, y*). In our simulations, we use Δ*t* = 0.96s, and with a resolution of 200 × 200 grid points, Δ*x* =1.5 *μ*m.

We further assess the stability of our numerical solutions by monitoring how the conclusions depend on the timestep Δ*t*. For this, we simulate a single cell-pair collision at different timesteps for both the velocity-aligning and front-front contact repolarization models. We repeat runs at each timestep 384 times and plot the average winning probability in Fig. 11. In particular, we note that the main simulation uses Δ*t* = 0.96s, which is at least a healthy factor of 2 smaller than a timestep at which convergence can become an issue.

**FIG. 11.**
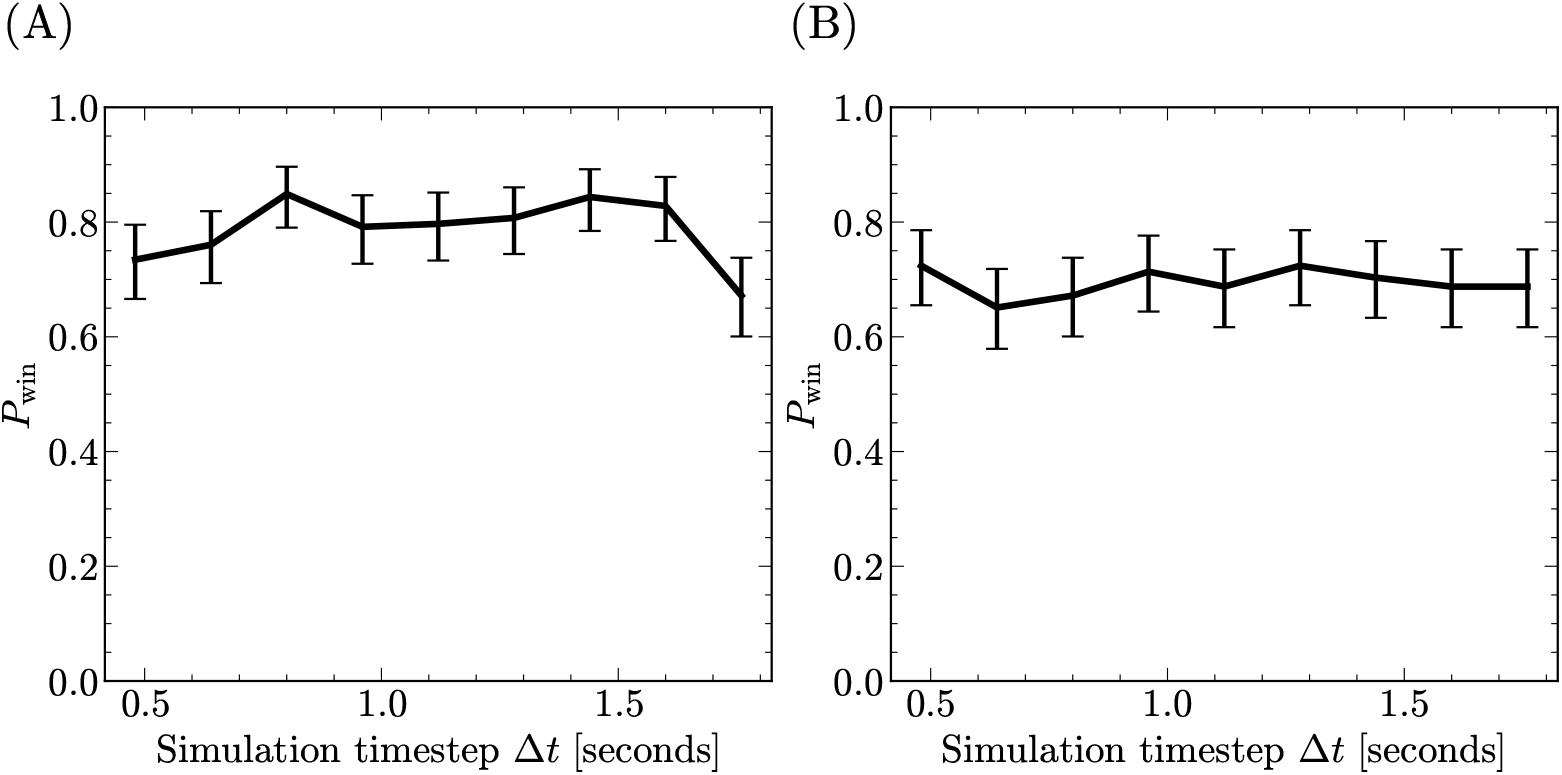
The average winning probability associated with a particular cell-pair collision is plotted against the timestep used in solving the equations of motion numerically. We see that the numerical solutions are stable over a healthy range of timesteps for both the (A) velocity-aligning and (B) front-front contact repolarization models. The averages are over *n* = 384 trials, and the error bars are the 95% binomial confidence intervals. Δ*t* = 0.96s is used in the main simulation.

## Appendix E: Solving a simplified contact repolarization model

We can solve a simplified version of the contact repolarization model analytically, determining the time it takes each cell to turn around. To do this, we neglect stochastic noise, as well as assuming that cell shapes (and hence the repolarization directions 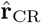) stay constant over the collision. We have defined cell polarity with an angle *ψ* associated with the vector 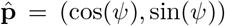, and contact repolarization by aligning the polarity of the cell toward the repolarization vector 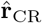 over a finite timescale, and we limit this interaction to only front-front contact. Mathematically, we describe this process with

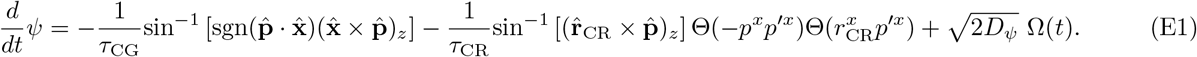

The first term is the contact guidance term ensuring the polarity of the cell remains parallel to the substrate. The second term models repolarization due to interactions with another cell with polarity 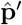, and the last term introduces Gaussian noise. 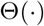 is the Heaviside step function and the two step functions here are used to limit contact repolarization to only happen when cells are in front-front contact.

Here, we consider the deterministic case with *D_ψ_* = 0 and solve Eq. (E1) analytically. Note that for unit vectors 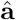 and 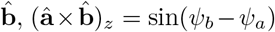, where 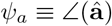 is the orientation of the unit vector 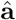, and similarly 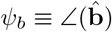. Now, consider a cell with polarity 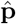 traveling to the right and about to collide head-on with another cell with polarity 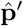 traveling to the left. Let *ψ* denote the direction of cell polarity, and 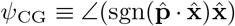 and 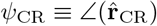 denote the contact guidance and contact repolarization angles, respectively. The contact guidance angle will be 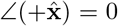 when *ψ* < *π*/2, then switch to 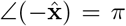 for *ψ* > π/2. Because the collision takes place between the cell fronts, 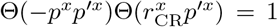. Lastly, we assume the repolarization vector 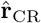 for the cell remains constant in time. Then, Eq. (E1) becomes

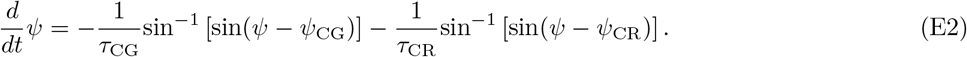

For Φ ∈ [−*π*/2,*π*/2], sin^−1^ [sin(Φ)] = Φ. However, given the initial condition *ψ*(*t* = 0) = 0 and *ψ*_CR_ > *π*/2, during the repolarization process, regions exist where Φ ≡ *ψ* – *ψ*_CR_ < −*π*/2 (Fig. 12: segment 1). For such cases, sin^−1^ [sin(Φ)] = −*π* – Φ. We can then rewrite Eq. (E2) piece-wise in terms of the value of *ψ*(*t*):

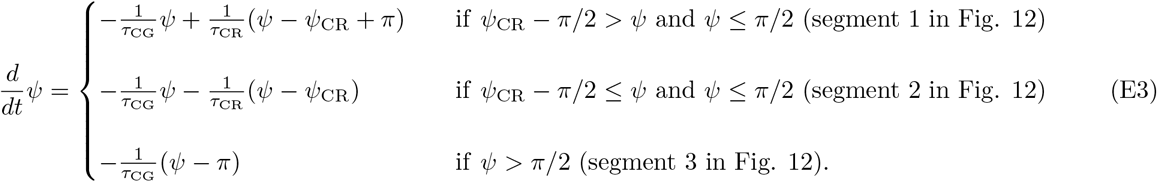

**FIG. 12.**
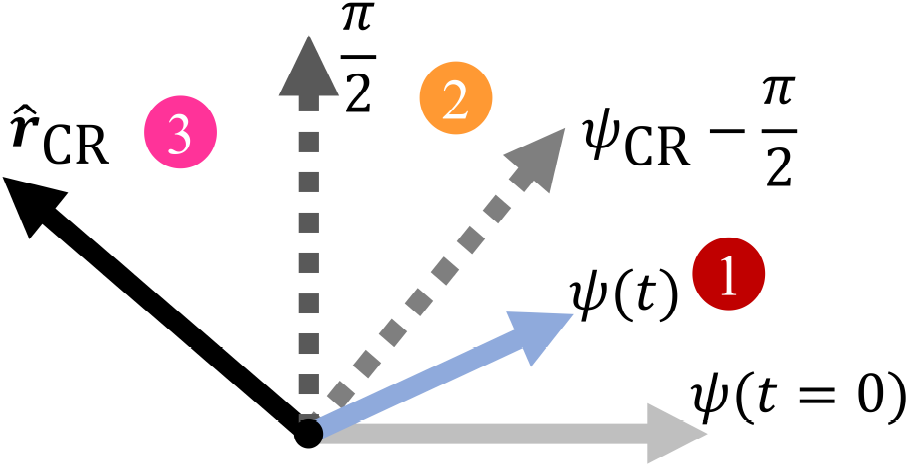
A schematic showing the time evolution of cell polarity *ψ*(*t*) within each of the three segments derived in Eq. (E3). Here, the cell starts at *ψ*_0_ = 0 and has a fixed repolarization vector 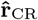. The color of each region refers to its analytical solution in Fig. 13A.

Note that when *ψ* > *π*/2, the cell has repolarized and its front is no longer in contact with the front of the other cell, and the repolarization term turns off – i.e. *τ*_CR_ → ∞ – and cell polarity relaxes toward *ψ*_CG_ = *π* with the time constant *τ*_CG_.

With a little algebra, we can write Eq. (E3) more clearly and reveal the effective timescales and steady-state angles for each segment:

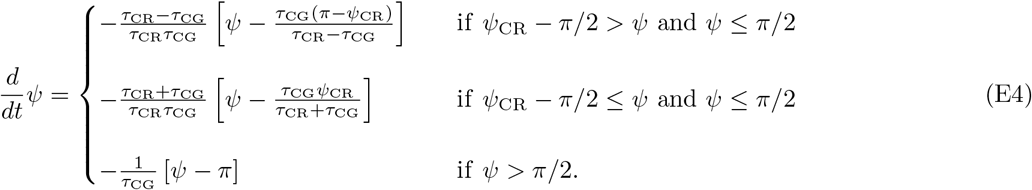

Each segment admits a solution of the form *ψ*_*i*_(*t*) = Φ_*i*_ + *c*_*i*_ exp[−*t*/*τ*_*i*_], where Φ_i_, *τ_i_*, and *c*_*i*_ are the steady-state angle, time constant, and constant of integration for each segment *i* = [1, 2, 3] (Fig. 12). Let us now focus on each segment individually:

1. When *ψ*_CR_ – *π*/2 > *ψ* and *ψ* ≤ *π*/2 (segment 1 in Fig. 12): Given *ψ*_1_(*t* = 0) = 0, we have *c*_1_ = Φ_1_. Thus,

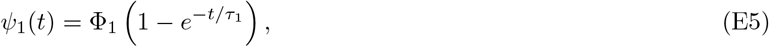

with 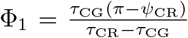 and 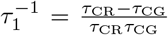. To ensure we proceed to the second region, we must have Φ_1_ ≥ *ψ*_CR_ – *π*/2. This condition translates to a constraint on the timescales:

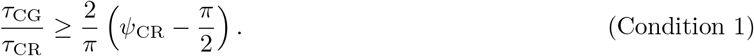 Lastly, we calculate the time at which the dynamics of cell polarity switch to that of the next segment – i.e. *t*′, such that *ψ*_1_(*t*′) = *ψ*_CR_ – *π*/2:

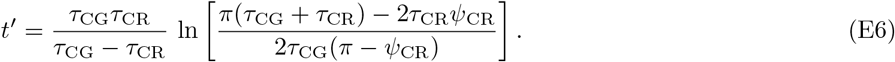
2. When *ψ*_CR_ – *π*/2 ≤ *ψ* and *ψ* ≤ *π*/2 (segment 2 in Fig. 12): The initial condition for this segment is *ψ*_2_(*t* = *t*′) = *ψ*_CR_ – *π*/2. Instead of plugging this directly into the exponential form and solving for *c*_2_, it would be easier to shift the solution temporally and let it begin at *t*′: *t* → *t* – *t*′. Then, *c*_2_ = *ψ*_CR_ – *π*/2 – Φ_2_, and

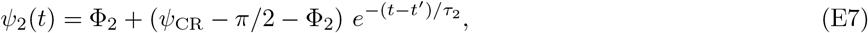

where 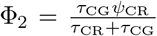 and 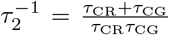. To ensure the cell can properly repolarize away from contact, we require Φ_2_ > *π*/2. This condition places another constraint on the timescales:

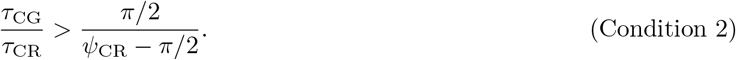 Lastly, we compute the time at which the dynamics of the cell polarity switch to that of the next segment – i.e. the repolarization time *t*″, such that *ψ*_2_(*t*″) = *π*/2:

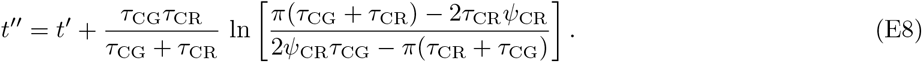
3. When *ψ* > *π*/2 (segment 3 in Fig. 12): The initial condition for this segment is *ψ*_3_(*t* = *t*″) = *π*/2. Again, if we shift this solution temporally by an amount *t*″, we can effortlessly compute the constant of integration, *c*_3_ = *π*/2 – Φ_3_. Then,

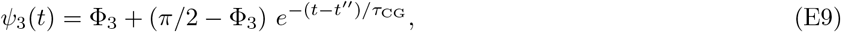

where Φ_3_ = *π*. Note that since the cell’s front is no longer in contact, the repolarization term turns off and cell polarity relaxes toward Φ_3_ with the time constant *τ*_CG_.

At last, we have found the full piece-wise solution to Eq. (E3):

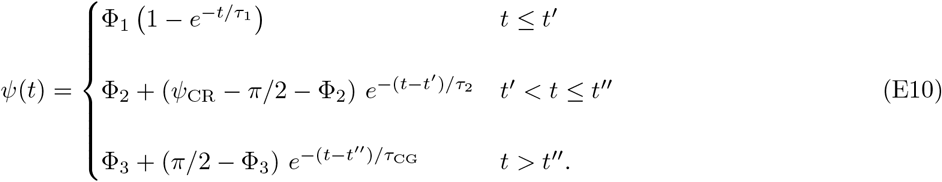

Here, *t*′ denotes the first transition point in time with *ψ*(*t*′) = *ψ*_CR_ – *π*/2, and *t*″ defines the time it takes the cell to repolarize, that is *ψ*(*t*″) = *π*/2. Fig. 13A plots these analytical solutions against the numerical solution of Eq. (E2).

**FIG. 13.**
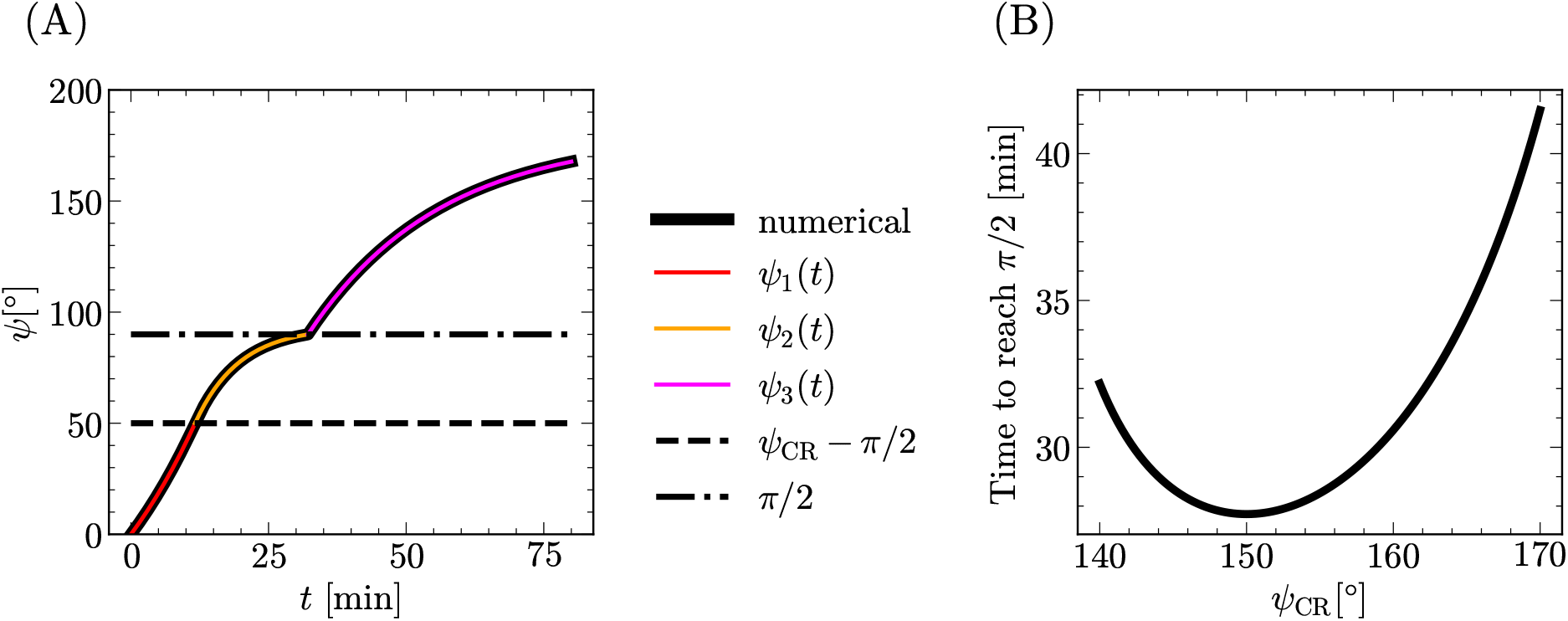
(A) Analytical solutions to the FFCR model are plotted for each segment against the numerical solution. Here, we consider a cell traveling to the right with *ψ*(*t* = 0) = 0, *τ*_CG_ = 2*τ*_CR_, and *ψ*_CR_ = 140°. Note that the angle *ψ*_3_(*t*) has not yet reached its steady state in this plot – it will asymptotically approach *π* at long times. (B) The time it takes cell polarity to reach *ψ* = *π*/2 starting from *ψ* = 0 (*t*″ in Eq. (E8)) is plotted as a function of the repolarization direction *ψ*_CR_.

The above derivation serves two main purposes. First, it allows us to choose the timescales of the model properly. As detailed above, Condition 1 and Condition 2 must be satisfied to ensure the polarity of the cell can trace through the appropriate angular range to undergo a proper repolarization. With some algebra, one can prove that Condition 2 is stricter than Condition 1. Thus, it suffices to satisfy the former. In our simulations, the roundest cells – which determine the lowest bound on the inequality – have a repolarization vector with *ψ*_CR_ ≈ 140° on average. Plugging this into Condition 2, we see that we must satisfy *τ*_CG_/τ_CR_ > 1.8 to ensure all cells deterministically repolarize. This restriction informed our choice of *τ*_CG_ = 2*τ*_CR_.

Moreover, the analytical solutions have allowed us to explicitly calculate the repolarization time for a cell whose polarity begins at *ψ* = 0. More precisely, this is the time it takes cell polarity to reach *ψ* = *π*/2, which is denoted by *t*″ in Eq. (E8). How does this repolarization time depend on the repolarization angle *ψ*_CR_? Consider the extreme – albeit unphysical – case of *ψ*_CR_ = *π*/2. Then, according to *ψ*2(*t*) from Eq. (E7), it would take an infinitely long time to reach *π*/2. As *ψ*_CR_ grows away from the vertical, cell polarity can reach *π*/2 more quickly. As an analogy, think of an overdamped harmonic oscillator that would take an infinitely long time to reach its equilibrium point. If a new equilibrium point is chosen past the previous one, then the former point can be reached in finite time. However, we do not expect *t*″ to decrease monotonically with increasing *ψ*_CR_. Consider another extreme – yet unphysical – case of *ψ*_CR_ = *π*. This puts us in the region where *ψ*_CR_ – *π*/2 > *ψ* and *ψ* ≤ *π*/2, and according to Eq. (E4), *d_ψ_*/dt = 0. This is indeed a point of unstable equilibrium for the cell’s polarity, exactly like a pendulum perfectly balanced to point “North” is stationary, but unstable. In this case, it would again take an infinitely long time to reach *π*/2. Thus, we would expect *t*″(*ψ*_CR_) to decrease from infinity as *ψ*_CR_ grows away from *π*/2, reach a minimum at an angle determined by the timescales, and grow toward infinity as *ψ*_CR_ nears *π*. The exact behavior of *t*″ is plotted in Fig. 13B for timescales used in simulation and for a subset of repolarization angles that were observed within our simulations, *ψ*_CR_ ∈ [140°, 170°].

What insight can we gain from *t*″(*ψ*_CR_)? Recall sin^−1^[sin(Φ)] = −*π* – Φ when Φ ∈ [−*π*, −*π*/2), which arises from the sawtooth nature of the function. In our simulations, the center-of-mass of flatter cells and the repolarization vector passing through it lie closer to the horizontal. Consequently, flatter cells have larger repolarization angles, and compared to rounder cells, they have an angular difference Φ ≡ *ψ* – *ψ*_CR_ that is more negative. Citing the sawtooth shape of sin^−1^ [sin(Φ)], we have argued that flatter cells take longer to repolarize as compared to rounder cells because the strength of their repolarization term is weaker. This has been our core justification for why flatter cells are more likely to win collisions under the front-front contact repolarization model. The repolarization time as a function of repolarization angle, which is plotted using simulation parameters, supports this notion to some extent: for *ψ*_CR_ > 150°, time to repolarize monotonically increases with increasing repolarization angle (Fig. 13B). This means that flatter cells take longer to flip, and when colliding with rounder cells, which repolarize sooner, they will emerge as the “winner”. *t*″(*ψ*_CR_) also tells us something very interesting: a flat cell with *ψ*_CR_ ≈ 150° will – deterministically – flip sooner and *lose* to a rounder cell with *ψ*_CR_ ≈ 140°. Do we have simulation points of this nature? Yes. Are they evidence against the notion that collision outcome correlates with how flat a cell is? That depends on how important the dip in *t*″(*ψ*_CR_) is. Our phase field simulations are *not* deterministic, but rather noisy. For two cells to have repolarization angles *ψ*_CR_ ∈ (140°, 150°), they would have to be very similar in their physical properties. In addition, the difference in the time to reach *π*/2 observed in Fig. 13B between *ψ*_CR_ = 140° and 150° is much smaller than that between 150° and 170°. Repolarization will also depend strongly on the initial polarity direction *ψ* at collision; when the angular diffusion coefficient *D_ψ_* is nonzero, this will vary stochastically. Which cell turns around first will also be affected by stochastic fluctuations in the polarity angle’s trajectory. Even though the flatter cell with *ψ*_CR_ = 150° would deterministically turn first and lose to the rounder cell with *ψ*_CR_ = 140°, angular noise will tend to wash out the small differences in repolarization time and drive the collision closer to a 50-50 outcome. This means that sufficiently large levels of noise can mask the influence of the dip in *t*″, and we would observe that flatter cells are generally slower to turn and more likely to win. However, if the system has low levels of noise, then it behaves more deterministically, and the dip in the repolarization time matters more. This is a potential reason for why relative contact angle – a measure of how flat a cell is – becomes a poor predictor of collision outcome when *D_ψ_* is very small (bottom row of Fig. 9 in main text).

## Appendix F: How accurately do *δv* and *δθ* predict *P*_win_?

One question motivating our work has been whether the outcome of collision can be predicted. We have shown that a single observable – relative speed *δv* or relative contact angle *δθ* – can robustly characterize the outcome of collision between cells with widely varying attributes. Given a value for one of these observables, how accurately can we predict the winning probability?

Tables II–V show two assessment metrics for each logistic regression performed on either of the predictors. A particular table (e.g. Table II) focuses on a given polarity mechanism – velocity-aligning (VA) or front-front contact repolarization (FFCR) – and angular diffusion coefficient *D_ψ_*. To compute a given assessment metric, we employ the *k*-fold cross-validation method. Here, we randomly shuffle and split the entire dataset obtained from simulations into *k* =10 equally sized segments. We then train the regression model on nine segments (*n*_train_ = 39, 917 points), and we compute the assessment metric on the remaining one segment (*n*_test_ = 4, 435 points). At the end, we have k = 10 assessment metrics, from which we report the average and standard deviation (Tables II–V).

**TABLE II.** VA model, *D_ψ_* = 0.075 rad^2^/hr

| Predictor | $\delta v$ | $\delta\theta$ |
| --- | --- | --- |
| Score | $79.5 \pm 0.8\%$ | $70.5 \pm 0.6\%$ |
| Brier Score | $0.140 \pm 0.003$ | $0.196 \pm 0.003$ |

**TABLE III.** VA model, *D_ψ_* = 0.0075 rad^2^/hr

| Predictor | $\delta v$ | $\delta\theta$ |
| --- | --- | --- |
| Score | $88.2 \pm 0.3\%$ | $73.8 \pm 0.4\%$ |
| Brier Score | $0.082 \pm 0.002$ | $0.172 \pm 0.002$ |

**TABLE IV.** FFCR model, *D_ψ_* = 0.075 rad^2^/hr

| Predictor | $\delta v$ | $\delta\theta$ |
| --- | --- | --- |
| Score | $64.3 \pm 0.7\%$ | $72.1 \pm 0.7\%$ |
| Brier Score | $0.222 \pm 0.002$ | $0.182 \pm 0.003$ |

**TABLE V.** FFCR model, *D_ψ_* = 0.0075 rad^2^/hr

| Predictor | $\delta v$ | $\delta\theta$ |
| --- | --- | --- |
| Score | $68.1 \pm 0.7\%$ | $84.2 \pm 0.3\%$ |
| Brier Score | $0.199 \pm 0.003$ | $0.107 \pm 0.002$ |

One widely used assessment metric is the score function, the percentage of correctly labeled points. Here, the class label of a point is determined by applying a binary threshold to the predicted winning probability: the class label of point *δv* is “win” if *P*_win_(*δv*) ≥ 0.5, else it is “loss”. Shown in the first row of Tables II–V, this metric yields a percentage value characterizing how successfully the model classifies observations. We find that when the velocity-aligning model is employed, relative cell speed predicts collision outcomes with 79.5 ± 0.8% (*D_ψ_* = 0.075 rad^2^/hr) success rate, and it is significantly better at predicting the winning probability compared to relative contact angle. Moreover, under the contact repolarization model, relative contact angle predicts collision outcomes with 72.1 ± 0.7% (*D_ψ_* = 0.075 rad^2^ /hr) success rate, and it is significantly the better predictor compared to relative speed. Additionally, as detailed in Section III D of the main text, as *D_ψ_* decreases, the winning probability tends toward a step-function, and we expect the score to increase. Comparing Tables III and V with Tables II and IV, we see a large increase in the score values of the appropriate predictor (*δv* for VA model, *δθ* for FFCR model).

We also compute the Brier Score, which is the mean squared error between the predicted *probability* and the class label (1: “win”, 0: “loss”) and avoids the binary threshold altogether. This metric is a cost function penalizing the classifier according to how incorrectly it labels points. As such, lower values correspond to better models. According to the Brier score, *δv* is the robust predictor of collision outcome under the VA model, while *δθ* best captures *P*_win_ under the FFCR model. Brier scores show that prediction is better (lower Brier score) at smaller values of *D_ψ_* for both models (Compare Tables III and V with Tables II and IV).

## Appendix G: Supplementary movies

Here we present movies of typical phase field collisions between two cells with different attributes. In each movie, the top panel tracks the evolution of the cells, while the bottom panels track the center-of-mass speed and contact angle of each cell as a function of time, respectively. Note that upon collision, we no longer compute these statistics and so the time series stop. In the following movies, the values of tension, adhesion to the substrate, and protrusion strength of the left cell are *γ* = 1.46*γ*_0_, *A* = 0.48*γ*_0_, and *β* = 5*γ*_0_, respectively. Meanwhile, the right cell has attributes *γ* = 1.1*γ*_0_, *A* = 0.4*γ*_0_, and *β* = 10*γ*_0_. These movies are mostly mean to be illustrative; note that the left cell’s parameters are not the default parameters. We have also set the angular diffusion coefficient to a small value, *D_ψ_* = 0.0075 rad^2^/hr so the collision is simpler to follow by eye.

- **Movie S1.** Collision of two cells whose polarity is modeled by the velocity-aligning (VA) mechanism with the alignment timescale *τ*_VA_ = 24 min.
- **Movie S2.** Collision of two cells whose polarity is modeled by the front-front contact repolarization (FFCR) mechanism with the alignment times *τ*_CG_ = 24 min and *τ*_CR_ = 12 min. When the cells approach each other head-on and form contact, the repolarization vector **r**_CR_ is computed according to Eq. 9 and plotted with a black arrow. Once one of the cells turns, head-head contact is lost and contact repolarization is turned off. The cells continue to travel as a train with their newly formed head-tail contact.

